# Phosphoinositides protonation states dictate AKT1 binding with membrane

**DOI:** 10.64898/2026.09.05.749365

**Authors:** Kirtika Jha, Meghadeepa Sarkar, Krishnakanth Baratam, Jayashree Nagesh, Thomas J. Pucadyil, Anand Srivastava

## Abstract

Phosphoinositides are key regulators of membrane-associated signaling, yet their electrostatic properties are commonly represented by a single nominal charge in molecular simulations and structural models. Using the essential protein kinase AKT1 as a model system, we show that phosphoinositide protonation microstates, both the degree and positional placement of protons, govern peripheral membrane-protein recognition. By integrating quantum-mechanics-derived headgroup parameters, microsecond-scale all-atom molecular dynamics, and umbrella-sampling free-energy calculations with the available solid-state NMR-derived lipid populations via Bayesian/Maximum Entropy reweighting, we account explicitly for the thermodynamic ensemble of phosphoinositide protonation states. This population-weighted free-energy framework provides a quantitative mechanistic rationale for the strict specificity of wild-type AKT1 toward PI(3, 4, 5)P_3_ and elucidates how the oncogenic sentry mutation (E17K) alters this specificity to enable high-affinity PIP_2_ binding. Furthermore, Proximity-based Labeling of Membrane-Associated Proteins (PLiMAP) assays experimentally confirm that AKT1 pleckstrin homology domain binding to distinct phosphoinositide species is differentially sensitive to pH. Together, our findings establish a direct connection between phosphoinositide protonation equilibria, binding energetics, and AKT1 lipid specificity. Beyond AKT1, these results demonstrate that the dynamic protonation microstates of anionic phospholipids act as a functional regulatory layer governing peripheral membrane-protein recruitment across diverse cellular microenvironments.

**Significance Statement:** In physiological conditions, AKT1 protein is highly specific to PI(3,4,5)P_3_ lipids and completely excludes PI(4,5)P_2_ lipids. However, a single charge reversal (oncogenic) mutation of sentry glutamate (E17K) on the membrane adaptor Pleckstrin Homology domain of AKT1 enhances its PIP_2_ binding remarkably. Our simulation data reveal that AKT1-PHD dissociation free-energies have remarkable variations with changes in lipid protonation state. By integrating our atomic simulation readouts with available ensemble-averaged low-resolution information from experiments using machine learning methods, we could explain the highly enhanced PIP_2_ specificity for the pathological mutant. The work highlights the degenerate roles of secondary messenger lipids as a function of membrane environments.

## Introduction

The essential PI3K/AKT/mTOR (phosphatidylinositol 3-kinase/ Protein Kinase-B/ mechanistic target of rapamycin) signaling pathway regulates cell survival, cell growth, differentiation, cellular metabolism, and cytoskeletal reorganization of cells. ^1–4^ One of the main proteins involved in this signaling pathway is AKT1 – a 57-kDa, 480 amino acids long Serine/Threonine kinase. The domain architecture of AKT1 includes the membrane-adaptor N-terminal pleckstrin homology domain (AKT1-PHD) followed by an unstructured linker region, a conserved kinase domain, and a flexible C-terminal regulatory domain. ^5–8^ The plasma membrane (PM) association of AKT1, which is a critical step in the pathway, is regulated by highly stereospecific binding of AKT1-PHD with the phosphatidylinositol-3,4,5-triphosphate (PIP_3_) lipids. Unlike most PH domains belonging to other proteins, such as GRP1, dynamin, PLC*δ*1, AKT1-PHD does not bind to PI(4,5)P_2_ but exhibits very high affinity for PIP_3_.^8–13^ Interestingly, a single charge reversal mutation of a sentry Glutamate residue to Lysine (E17K) is known to increase the protein’s affinity for PI(4,5)P_2_ lipids.^14,15^

PIP or Phosphoinositides (diacyl-phosphatidylinositol phosphates) lipids are quite different from constitutional phospholipids and form only a minor pool of membrane lipids. However, they are important second messengers and play a critical role in a broad range of signaling events.^16^ PIP lipids have a glycerol backbone esterified to two fatty acid chains and phosphate and attached to a polar head group, which is the cyclic polyol myoinositol (CHOH)_6_. This inositol head group has free hydroxyl groups at positions C2 through C6 that can be differentially phosphorylated by cytoplasmic lipid kinases at positions C3, C4, and C5. As a result, phosphoinositide lipid species can naturally occur in seven combinatorial phosphorylated forms, some of which often behave as distinct organelle membrane markers.^17,18^ Among the seven naturally occurring phosphorylated forms, phosphatidylinositol-3,4,5-trisphosphate (PIP_3_) bears the maximum charge in its headgroup and localizes at the plasma membrane along with the phosphatidylinositol-4,5-diphosphate (PIP_2_) species. Structurally, PIP_3_ and PIP_2_ only differ by a phosphate group, resulting in different charges and stereochemistry that dictate their specificity to membrane adaptors such as PH domain. Notably, these PIPs can exist in multiple protonation states, such as fully protonated, partially protonated, or deprotonated, based on their local environment conditions (e.g., pH, neighboring lipids, and the concentration of ions).

Detailed solid-state NMR studies from Koojiman and co-workers^19^ have shown that the protonation states of PIP lipids follow a complex trend. The work also emphasized that a change in pH not only alters the overall PIP charge but also changes the distribution of the charge within the headgroup. In the past, related work by Arne Gericke and co-workers had also studied the effect on the protonation of PIP_2_ in the presence of the neighboring hydrogen-bond donor lipids such as phosphatidylethanolamine (PE) and phosphatidylinositol (PI).^20^ They were able to show that these neighboring lipids not only influenced the charge of PIP_2_ via hydrogen-bond formation with its phosphomonoester groups but also the lateral distribution of PIP_2_ was affected in the case of PI, resulting in the formation of nanoscale domains rich in PIP lipids.^19^ The membrane micro-environment dictates the relative populations of the ionization/protonation states of these lipids, which in turn affects the membrane association behavior of peripheral membrane proteins. However, it is non-trivial to accurately obtain the relative populations of the protonation states of the lipids due to the ensemble-averaged low-resolution binding-assay readouts from the experiments.

It is extremely challenging experimentally to investigate such a dynamic microenvironment and obtain accurate insights into all possible different protonation states. Molecular dynamics (MD) simulations can be used as a complementary method to model and probe the various possibilities arising in such systems. In this work, we use atomic-scale MD simulations and umbrella sampling-based free energy calculations to understand the molecular mechanism involved in AKT-PIP interaction depending on the protonation state of PIP lipids. Additionally, we also explored the molecular-level details responsible for the loss of PIP_3_ specificity due to the charge reversal mutation (E17K). In particular, we explore the effect of protonation states of PIP lipids on binding geometries and the free energy of dissociation. For a given charge state, we also explore the effect of proton position on the lipid headgroup to get insights into the association and dissociation dynamics of the AKT1 PH domain. PIP_3_ has eight distinct protonation states with one fully protonated state, three singly deprotonated states, three doubly deprotonated states, and one state with full deprotonation.^21^ Similarly, PIP_2_ has four distinct protonation states. All different headgroups of PIP_3_ and PIP_2_ considered in this study have been shown in **Figure 1A**.

**Figure 1:**
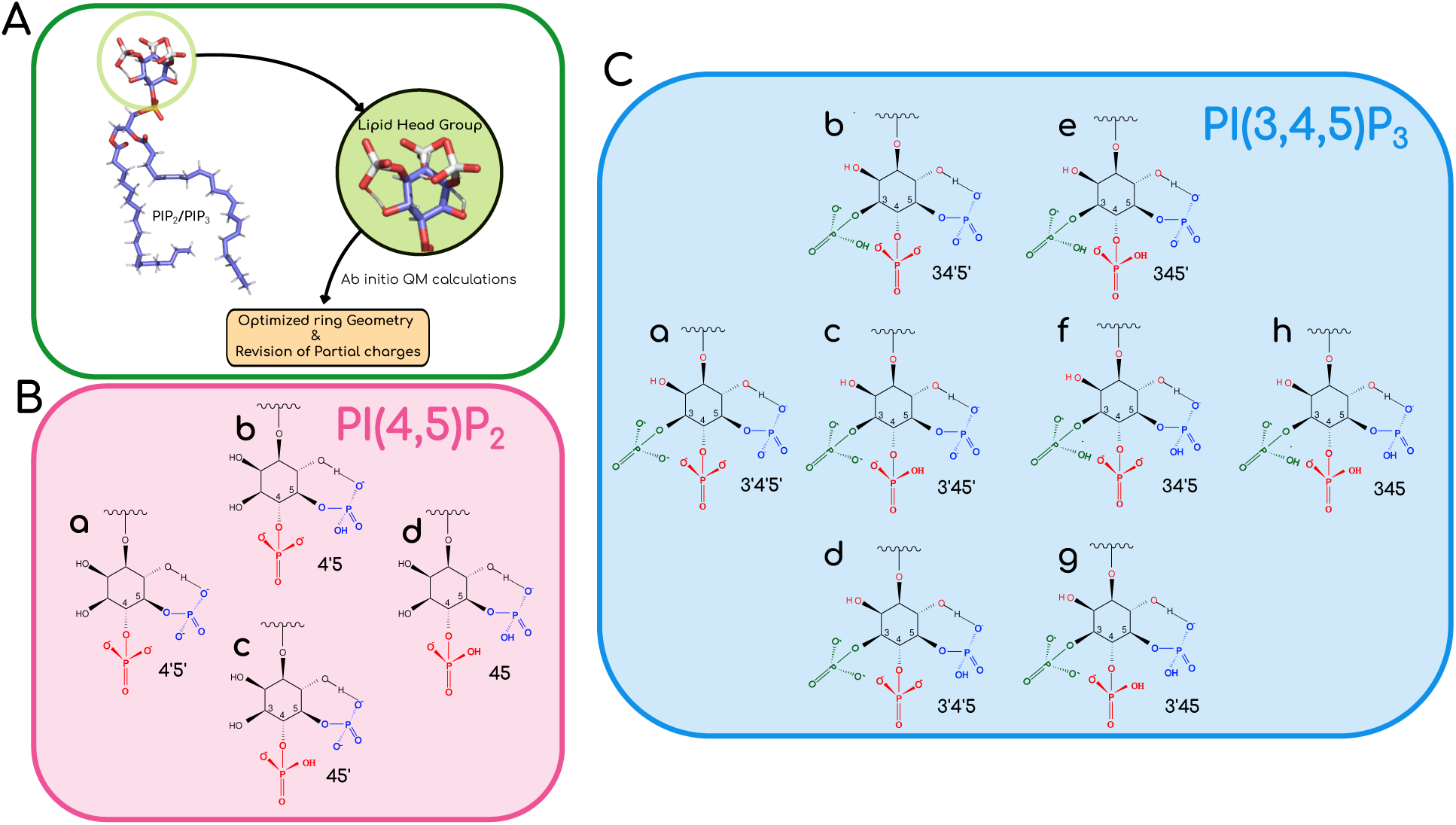
(A) Calculation of partial charges of differently protonated lipid headgroups using Ab initio quantum calculation. (B) Protonation states of PIP_2_ (a) PI(4’,5’)P_2_(No protonation) (b) PI(4’,5)P_2_ (c) PI(4,5’)P_2_ (Single protonation) (d) PI(4,5)P_2_ (Double protonation) (C) Protonation states of PIP_3_ (a) PI(3’,4’,5’)P_3_ (No protonation) (b) PI(3,4’,5’)P_3_ (c) PI(3’,4,5’)P_3_ (d) PI(3’,4’,5)P_3_ (Single protonation) (e) PI(3,4,5’)P_3_ (f) PI(3,4’,5)P_3_ (g) PI(3,4’,5’)P_3_ (Double protonation) (h) PI(3,4,5)P_3_ (Triple protonation)

The remainder of this article is structured as follows. This Introduction section is followed by the Materials and Methods section, where we list the details of *ab initio* Quantum Mechanics based electrostatics calculations to extract the altered partial charge distribution and optimized geometries for the PIP_2_ and PIP_3_ head group for a given protonation state and use the information in our all-atom MD simulations. We also provide details about the various all-atom molecular dynamics systems considered for simulations in this work, along with the details for the umbrella sampling calculations for all the systems under consideration. Following the simulations, we discuss the methodological details of how we have used the Maximum Entropy algorithm^22,23^ to integrate our simulation results with ssNMR data on lipid fractional populations^19^ and biochemical binding affinity experimental data,^15^ which together provide us with additional insights into the bilayer-AKT interactions. The Materials and Methods section is followed by the Results and Discussion section, where we report our findings and highlight the role of lipid protonation on binding affinities of peripheral membrane proteins. We find that the binding geometries and dissociation free energies have remarkable variations with protonation states of the lipids. The insights gained at the binding interfaces from the molecular simulations and the free-energy calculations were used to explain the equilibrium parameters data reported earlier from biophysical experiments with AKT1 PHD.^15^ We summarize our work in the Conclusion section. All input files needed to carry out the *ab initio* calculations and molecular simulations are made available on the following Zenodo repository: https://zenodo.org/uploads/17230768. All codes and raw data related to Maximum Entropy calculations are also provided in the same repository. We have also deposited the trajectories and some related movie files in the same repository.

## Materials and Methods

### *Ab initio* calculation of partial charges for headgroups with optimized geometry

In this work, we chose the Hu-Lu-Yang (HLY) scheme, where partial charges are obtained by minimizing the difference between the electrostatic potentials from Quantum Mechanics (QM) calculation and from atomic multipoles. The difference between this scheme and other ESP-fitting schemes is that the objective function is rotationally invariant to molecular orientation and varies smoothly with changes in molecular geometry. ^24^ All calculations are carried out in the Gaussian version G09-E01. We use B3LYP/6-311++G(d,p) for all-atom geometry optimization and charge calculation, with a polarizable continuum model for the water solvent. By this means, the partial charges for all possible headgroups of PIP_2_ and PIP_3_ were calculated. The schematic for the workflow is shown in **Figure 1A**.

### Incorporation of differentially protonated PIP**_2_**/PIP**_3_** lipids in force field

After calculating charges in the headgroup atoms for different protonation states, we replaced the existing headgroup in the full lipid with the newly generated lipid headgroups. Only rotational and translational modes were allowed for the superimposition of the headgroups. The charges were mapped onto the head groups as shown in **Supplemental Tables S1 and S2**. Further, these full PIP lipids with differentially protonated headgroups need to be added to the existing forcefield. The topology information for these lipids, along with the charges, was added in the required files of the forcefield. The new forcefield files for all the protonation states are available in the Zenodo repository.

To uniquely represent the different protonation states of the PIP_2_ and PIP_3_ lipids, we follow the notation such that an apostrophe on the number in the bracket indicates deprotonated PO_4_ on that location. For example, PI(4’,5’)P_2_ stands for fully deprotonated PIP_2_ and PI(4,5)P_2_ stands for full protonated PIP_2_ lipid. PI(4’,5)P_2_ stands for PIP_2_ with PO_4_ on the fourth carbon in the ring as deprotonated and the fifth one as protonated. The same notation is carried over for PIP_3_ lipids as well.

### All-Atom Simulations

We performed a set of 24 all-atom molecular dynamics (AAMD) simulations with both WT and mutant (E17K) AKT1 PHD on a bilayer to understand the dynamics and the effect of differently protonated PIP on protein binding. The initial orientation of AKT1-PHD on the bilayer was chosen based on the PDB ID:1H10, the solved crystal structure of AKT1 PHD bound with 4IP ligand. ^25^ All simulations were set up in such a way that the PHD was placed around 8-10 Å away from the bilayer surface. The simulations evolved over time until a bilayer-bound configuration via PIP headgroup was attained. Each protein-bilayer system comprised of PHD and 100 lipids per leaflet and with bilayer composition in the ratio of 80:19:1 (PC:PS:PIP). The systems were solvated and ionized with 150 mM KCl. Each of these systems was energy minimized and equilibrated for 50 ns in both NVT and NPT ensembles prior to setting up for a production run ranging from 500 to 1000 ns per system. We used CHARMM36m force-field^26^ for protein and CHARMM36 force-field^27^ for lipids. Non-bonded forces were calculated with a 12 Å cut-off (10 Å switching distance). Long-range electrostatic forces were calculated at every other time step using the Particle mesh Ewald method.^28^ The system was maintained at a temperature of 310 K and pressure of 1 atm using a Nose–Hoover thermostat^29^ and Parrinello–Rahman (with semi-isotropic coupling) barostat^30^ with time constants 1.0 and 5.0 *ps^−^*^1^, respectively. A set of 12 WT and 12 mutant protein-bilayer systems were simulated with a total simulation time of *∼* 15 *µ*s as shown in **Table 1**.

**Table 1:** Details of system set up for AAMD simulations with WT and E17K AKT1 PHD along with their protonation sites, charges and their length of simulation.

| Bilayer Composition | PIP lipid | Charge | Protonation sites | WT | E17K |
| --- | --- | --- | --- | --- | --- |
| POPC:POPS:PIP<sub>3</sub> | PI(3',4',5')P<sub>3</sub> | -7 | None | 1 $\mu s$ | 0.5 $\mu s$ |
| POPC:POPS:PIP<sub>3</sub> | PI(3,4',5')P<sub>3</sub> | -6 | P3 | 1 $\mu s$ | 0.5 $\mu s$ |
| POPC:POPS:PIP<sub>3</sub> | PI(3',4,5')P<sub>3</sub> | -6 | P4 | 1 $\mu s$ | 0.5 $\mu s$ |
| POPC:POPS:PIP<sub>3</sub> | PI(3',4',5)P<sub>3</sub> | -6 | P5 | 1 $\mu s$ | 0.5 $\mu s$ |
| POPC:POPS:PIP<sub>3</sub> | PI(3,4,5')P<sub>3</sub> | -5 | P3,P4 | 1 $\mu s$ | 0.5 $\mu s$ |
| POPC:POPS:PIP<sub>3</sub> | PI(3,4',5)P<sub>3</sub> | -5 | P3,P5 | 1 $\mu s$ | 0.5 $\mu s$ |
| POPC:POPS:PIP<sub>3</sub> | PI(3',4,5)P<sub>3</sub> | -5 | P4,P5 | 1 $\mu s$ | 0.5 $\mu s$ |
| POPC:POPS:PIP<sub>3</sub> | PI(3,4,5)P<sub>3</sub> | -4 | P3,P4,P5 | 1 $\mu s$ | 0.5 $\mu s$ |
| POPC:POPS:PIP<sub>2</sub> | PI(4',5')P<sub>2</sub> | -5 | None | 0.6 $\mu s$ | 0.5 $\mu s$ |
| POPC:POPS:PIP<sub>2</sub> | PI(4,5')P<sub>2</sub> | -4 | P4 | 0.6 $\mu s$ | 0.5 $\mu s$ |
| POPC:POPS:PIP<sub>2</sub> | PI(4',5)P<sub>2</sub> | -4 | P5 | 0.6 $\mu s$ | 0.5 $\mu s$ |
| POPC:POPS:PIP<sub>2</sub> | PI(4,5)P<sub>2</sub> | -3 | P4,P5 | 0.6 $\mu s$ | 0.5 $\mu s$ |

### Umbrella sampling simulations

To investigate the protonation-dependent conformational dynamics and to calculate the associated dissociation-free energy profiles, we performed umbrella sampling molecular dynamics (US-MD) simulations.^31^ The bound configuration obtained from AAMD simulation was used as the starting configuration for setting up the first window of US-MD simulation. Further, windows were set up by shifting the position of the bound PHD by placing it away from the membrane surface at an interval of 1 Å up to a distance of 25 Å as shown in **Figure 2**.

**Figure 2:**
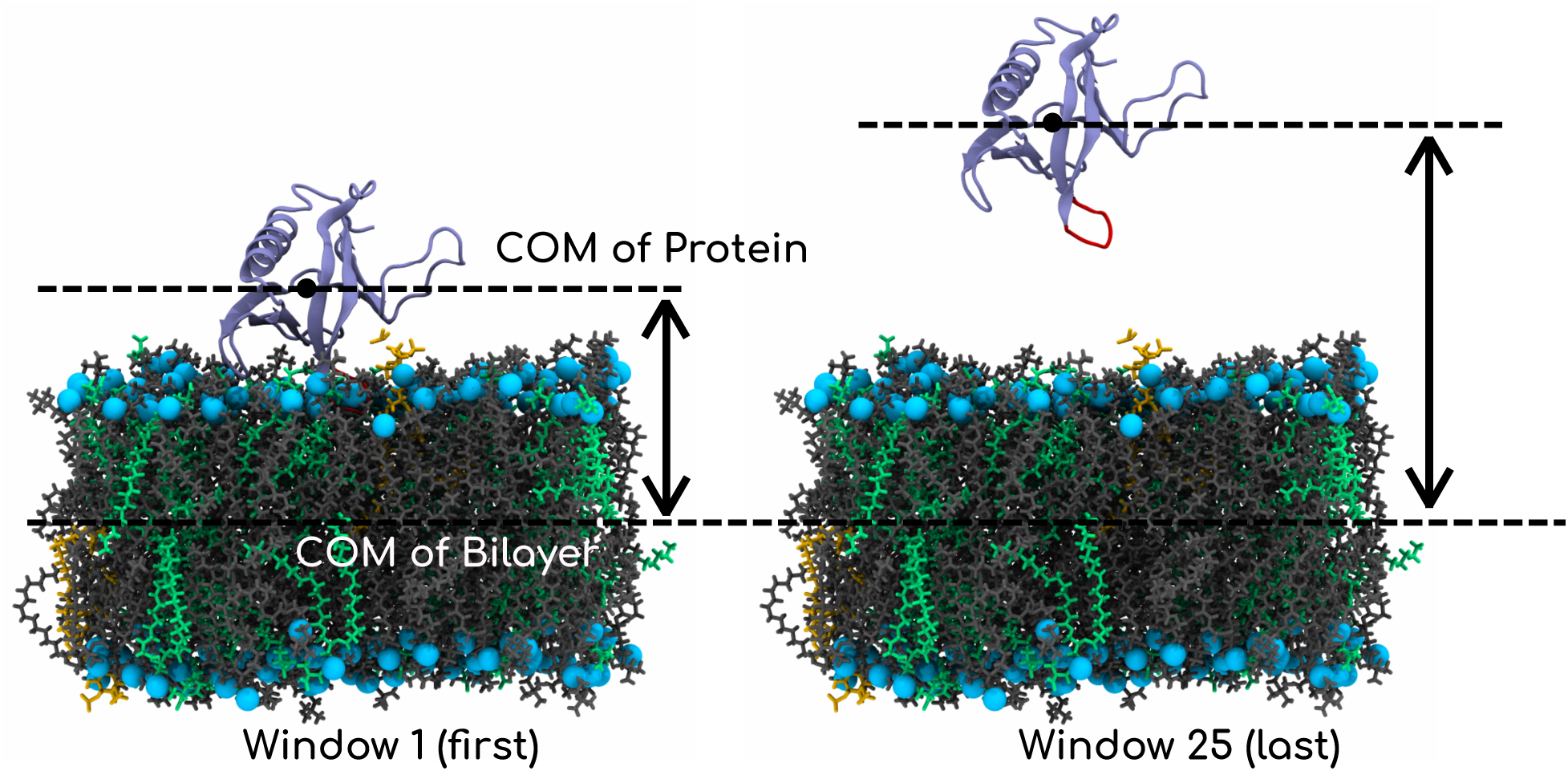
Snapshot of first and last window used for umbrella sampling simulation using distance between the COM of the membrane and the COM of protein (PHD) as the reaction coordinate. PC and PS lipids are shown in grey and purple lines respectively. PIP lipids are shown in vdW representation. AKT1-PHD is shown in cartoon representation. Water and ions are not shown for the clarity of image.

Umbrella sampling calculations were performed with a minimum distance between the center of mass of the PHD and the membrane midplane as the reaction coordinate. In total, 25 windows were generated, each simulated for 50 ns with an accumulated sampling of 1.25 *µ*s. In all these windows, we used a force constant (k) of 1000 kJ/mol/nm^2^. The potential of mean force (PMF) profile was then calculated using the gmx wham tool.^32^ The error along the PMF profile was obtained by performing a bootstrap analysis with 100 bootstraps. All PMF profiles were shifted to maintain the PMF value at the membrane plane at 0 kcal/mol value for reference. The same protocol was used for generating the PMF profiles of all the systems of interest.

### Functional forms of fraction of individual protonation states of PIP**_3_** and PIP**_2_**

Here, we elucidate the prescriptions that we apply to obtain the fraction of individual protonation states of PIP_3_ and PIP_2_ as a function of pH, along with the associated uncertainty. A detailed description of the experimental data and methods used to obtain the parameters in the following equations can be found in the original work of Koojiman and co-workers.^19^ Here, we represent the concentrations of individual states of PIP_3_ using the notation [345], [34′5], [345′], [3′45], [34′5′], [3′45′], [3′4′5] and [3′4′5′]. The sum of concentrations of individual protonation states of PIP_3_ is denoted by [*PIP*_3_]. The fraction of each protonation state is given by *f*_3′45_ = [3′45]*/*[*PIP*_3_], *f*_34′5_ = [34′5]*/*[*PIP*_3_], *f*_345′_ = [345′]*/*[*P IP*_3_], *f*_3′4′5_ = [3′4′5]*/*[*PIP*_3_], *f*_3′45′_ = [3′45′]*/*[*P IP*_3_], *f*_34′5′_ = [34′5′]*/*[*P IP*_3_], *f*_3′4′5′_ = [3′4′5′]*/*[*P IP*_3_], and *f*_345_ = 1 *−* (*f*_3′45_ + *f*_34′5_ + *f*_345′_ + *f*_3′4′5_ + *f*_3′45′_ + *f*_34′5′_ + *f*_3′4′5′_).

The rates of interconversion among the different protonation states are denoted either by the dissociation constants or equilibrium constants. *pK_a_*_1_, *pK_a_*_2_, and *pK_a_*_3_ denote the negative logarithm of dissociation constants associated with deprotonation of the fully protonated state ([345]) to singly deprotonated states [3’45], [34’5] and [345’], respectively. The constants *pK_a_*_7_ and *pK_a_*_8_, pertain to the deprotonation reactions of [3’45] to [3’4’5] and [3’45’] respectively. *pK_a_*_9_ and *pK_a_*_10_, pertain to the deprotonation reactions of [34’5] to [3’4’5] and [34’5’] respectively. *pK_a_*_11_ and *pK_a_*_12_, pertain to the deprotonation reactions of [345’] to [3’45’] and [34’5’] respectively. *pK_a_*_16_, *pK_a_*_17_,and *pK_a_*_18_, pertain to the deprotonation reactions of [345’], [3’45’] and [34’5’] to the fully deprotonated state [3’4’5’]. *K*_4_, *K*_5_, and *K*_6_ are the equilibrium constants used to describe the interconversion among singly deprotonated states while *K*_13_, *K*_14_, and *K*_15_ describe the exchange among doubly deprotonated states.

Using the Henderson-Hasselbalch^33^ equation on individual protonation states and eliminating the interconversion equilibrium constants, we arrive at the set of equations (Equations and 2) that allow us to evaluate the fraction of each protonation state as a function of pH, thus reproducing the observations made by Kooijiman and co-workers.^19^ We further quantified the uncertainty of each of these fractions using the error reported on the *pK_a_*s. We represent the fraction of each protonation state as a function of pH in **Figure 3**.

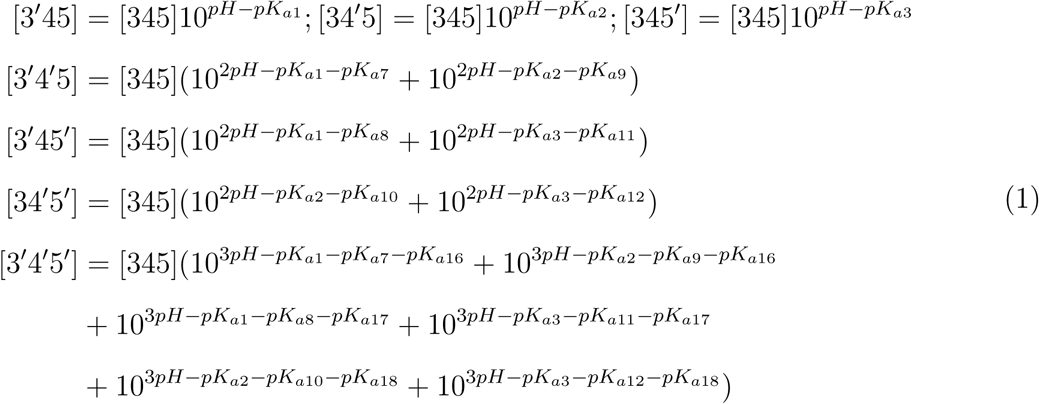

Similarly, the concentrations of individual protonation states of PIP_2_ are represented using [45], [4′5], [45′] and [4′5′]. The sum of the concentrations of individual protonation states of PIP_2_ is denoted by [PIP_2_]. *pK_a_*_1_, and *pK_a_*_2_ denote the negative logarithm of dissociation constants associated with deprotonation of fully protonated state([45]) to singly protonated states [4’5] and [45’] respectively. *pK_a_*_4_ and *pK_a_*_5_ are the constants associated with the conversion of these singly protonated states to the fully deprotonated state ([4’5’]). *K*_3_ is the equilibrium constant to describe the interconversion among singly deprotonated states. Upon similar transformation as done with PIP_3_ using Henderson-Hasselbalch^33^ equation on individual protonation states and eliminating the interconversion equilibrium constant, we get the fraction of each protonation state *f*_4′5_ = [4′5]*/*[*PIP*_2_], *f*_45′_ = [45′]*/*[*P IP*_2_], *f*_4′5′_ = [4′5′]*/*[*P IP*_2_], and *f*_45_ = 1 *−* (*f*_4′5_ + *f*_45′_ + *f*_4′5′_)

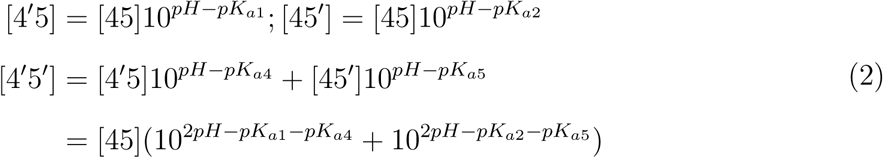

**Figure 3:**
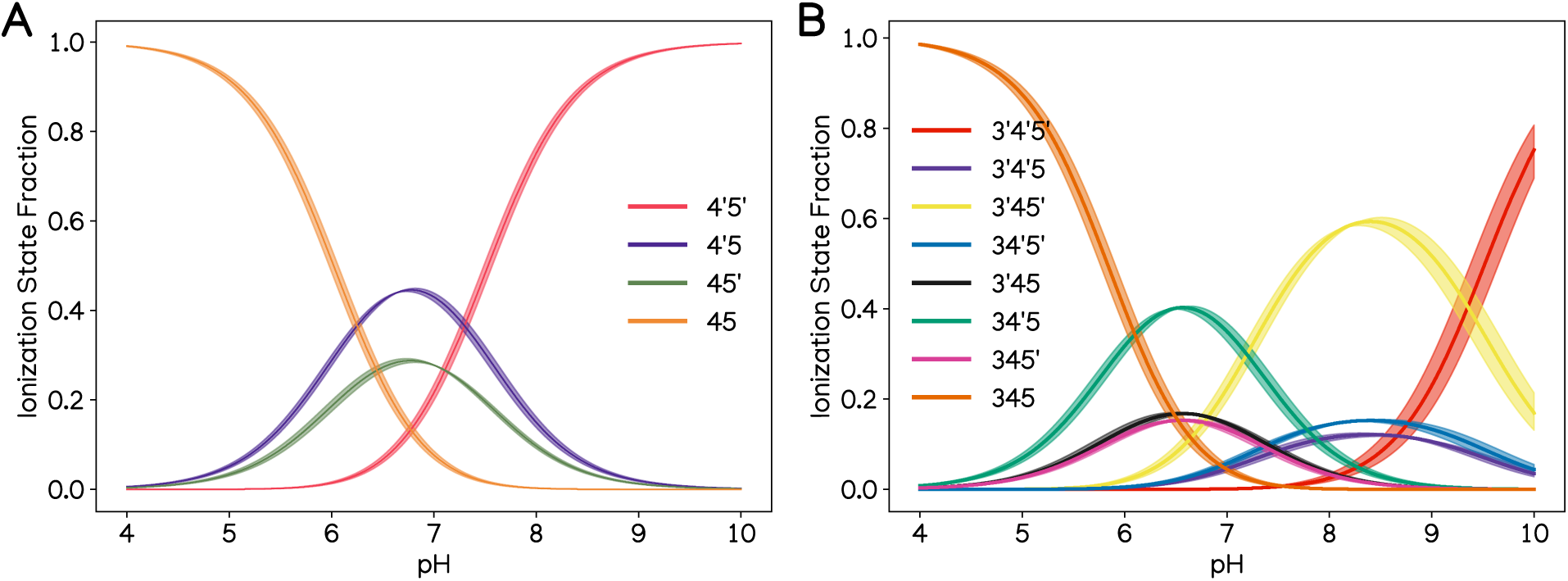
Fraction of each protonation state of (A) PIP_2_ and (B) PIP_3_ as a function of pH reproduced using constants along with their associated uncertainty from ssNMR-based study^19^

### Proximity-Based Labeling of Membrane-Associated Proteins (PLiMAP)

The PLiMAP assay was performed as previously described. ^34,35^ Briefly, purified proteins and vesicles were mixed at a 1:100 protein-to-lipid molar ratio in the designated pH buffers and incubated for 30 min at room temperature in the dark. For photo-crosslinking, the reaction mixtures were exposed to 365 nm UV light in a UVP CL-1000L crosslinker at an energy density of 200 mJ/cm² for 1 min. Samples were subsequently resolved by 15% SDS-PAGE, and the protein-associated BDPE fluorescence was captured using a Typhoon Biomolecular Imager (Amersham). Following fluorescence imaging, the gels were stained with Coomassie Brilliant Blue (CBB) and imaged using an iBright 1500 Imaging System (Invitrogen). Protein-lipid binding was quantified by defining a region of interest (ROI) around each protein band using image analysis. The local background intensity was subtracted from the band intensity, and the resulting net fluorescence values were plotted. Gels with WT and the E17K mutant were imaged together. Fluorescence was normalized to the sample showing maximum binding, which was with WT on PI(3,4,5)P3 at pH 6.0. Images were analyzed using Fiji,^36^ and data were plotted and analyzed using GraphPad Prism (Version 8.4.3).

### DNA constructs for PLiMAP assay

The AKT1 PH domain was amplified from Addgene plasmid #51465 (a gift from Dr. Tamas Balla, Budapest, Hungary) and cloned into a pET15b expression vector. The resulting construct encoded the protein with an N-terminal 6xHis tag and a C-terminal StrepII tag. The oncogenic E17K mutation was subsequently introduced via site-directed PCR mutagenesis. The sequences of all constructs were verified and confirmed by DNA sequencing.

### Protein expression and purification for PLiMAP assay

Recombinant proteins were expressed in *Escherichia coli.* T7 Express cells (New England Biolabs). Bacterial transformants were grown at 37 °C to an optical density (OD) of 0.6-0.8, followed by induction with 0.5 mM IPTG overnight at 18°C. Cultures were pelleted by centrifugation and stored at −40°C. For purification, frozen cell pellets were thawed in lysis buffer containing 20 mM HEPES (pH 7.4), 500 mM NaCl, 1% Triton X-100, and 1 mM phenylmethylsulfonyl fluoride (PMSF), and then disrupted via sonication. The lysate was clarified by centrifugation at 30,000g for 20 min at 4°C. The resulting supernatant was incubated with TALON Metal Affinity Resin (Takara Bio). The resin was washed with 20 mM HEPES (pH 7.4) and 500 mM NaCl, and the bound protein was eluted using the same buffer supplemented with 50 mM EDTA. The eluate was subsequently loaded onto a 5 mL StrepTrap HP column (GE Healthcare). After washing the column with 20 mM HEPES (pH 7.4) and 500 mM NaCl, the resin was equilibrated with a low-salt buffer consisting of 20 mM HEPES (pH 7.4) and 150 mM NaCl. Finally, the target protein was eluted using 2.5 mM desthiobiotin in the same low-salt buffer. Purified proteins were dialyzed against four distinct assay buffers, each containing 150 mM NaCl: 20 mM HEPES at pH 6.0, 7.0, or 8.0, and 20 mM Tris at pH 9.0. Following dialysis, the samples were centrifuged at 100,000g for 30 min at 4°C to remove any protein aggregates prior to use in downstream assays.

### Vesicle preparation for PLiMAP assay

1,2-dioleoyl-sn-glycero-3-phosphocholine (DOPC), 1,2-dioleoyl-sn-glycero-3-phospho-(1’-myoinositol-4’,5’-bisphosphate) (ammonium salt) (PI(4,5)P2), and 1,2-dioleoyl-sn-glycero-3-phospho(1’-myo-inositol-3’,4’,5’-trisphosphate) (ammonium salt) (PI(3,4,5)P3) were purchased from Avanti Polar Lipids. The UV-activable, diazirine-containing fluorescent lipid probe BODIPY-diazirine phosphatidylethanolamine (BDPE) was synthesized as previously described.^34,35^ Lipids were blended in glass tubes at a 98:1:1 molar ratio of DOPC to phosphoinositide (PI(4,5)P2 or PI(3,4,5)P3) to BDPE and dried under a high vacuum for 30 min to form a thin film. The dried lipid films were then hydrated with deionized water to a final total lipid concentration of 1 mM. Following hydration at 50°C for 30 min, the lipid suspensions were vigorously vortexed and sequentially extruded through 100 nm pore-size polycarbonate filters (Whatman) to generate large unilamellar vesicles (LUVs).

## Results and Discussions

### AKT1-PHD exhibit facile membrane association with all protonation states of PIP**_2_** and PIP**_3_** through Variable Loop 1

In our study, we set up and simulated 24 different PHD-bilayer systems. Here, we report membrane association data for bilayers containing eight possible differently protonated PIP_3_ lipid and four different PIP_2_ lipid states with both WT and mutant PHD, as mentioned in **Table 1**. Over the course of the simulation, we tracked the "z-distance" of each residue on the AKT-PHD from the phosphate plane of the bilayer in the z-direction. A representative bound state is shown in **Figure 4A**. In **Figure 4B and 4C**, we report the consolidated average z-distance observed over the last 50 ns of the trajectory for PIP_3_ and PIP_2_ states, respectively. A z-distance value *<* 0 indicates the insertion of the protein residue into the membrane. All 24 systems in WT and mutant states show some degree of negative values in the variable loop 1 (VL1) region of the AKT1-PHD.

**Figure 4:**
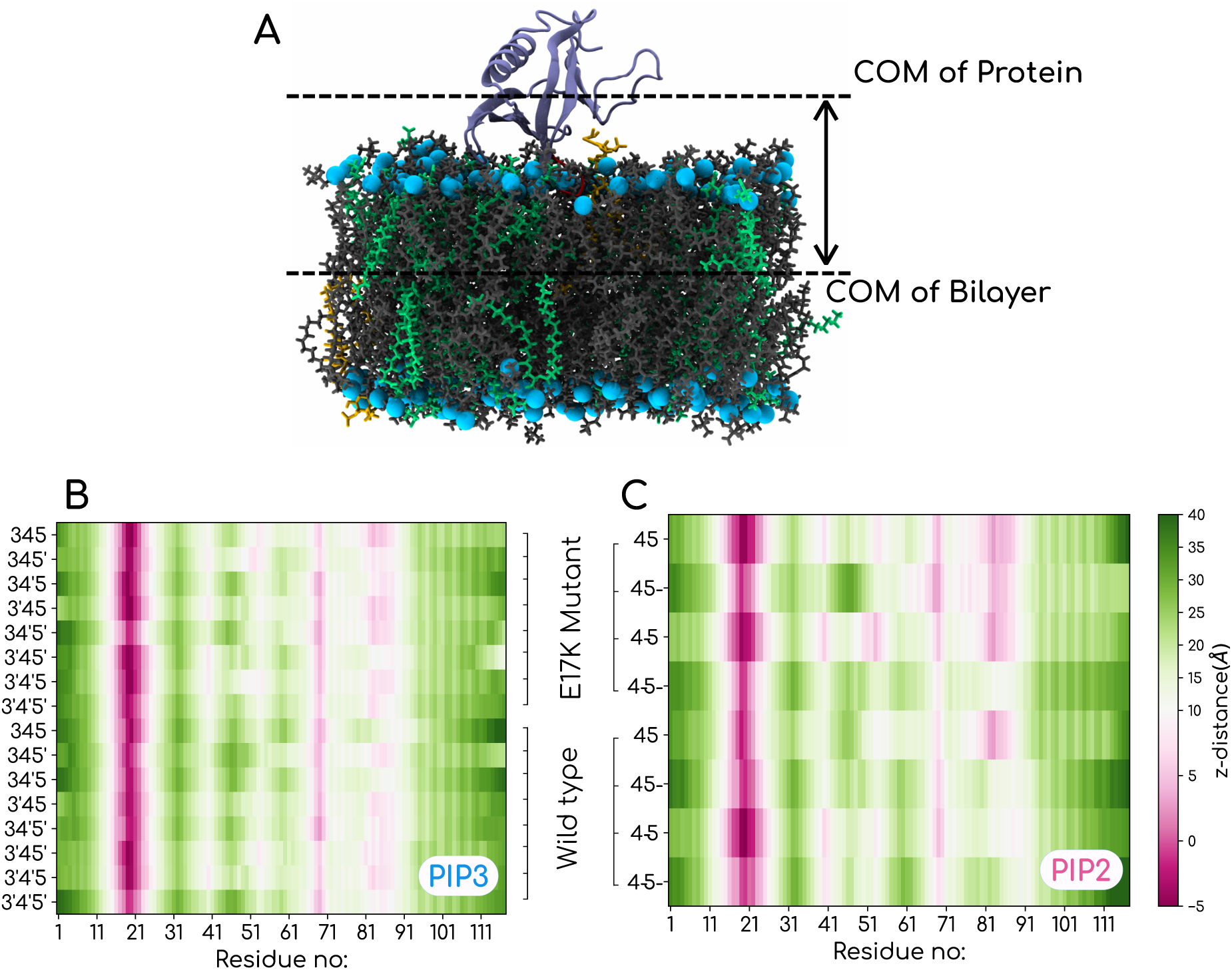
A) Representative image of an AKT1-PHD bound to PIP lipid. B) Heatmap showing z-distance of each residue of AKT1 PHD bound state from the bilayer plane for both wildtype and mutant PIP_3_ states and C)PIP_2_ states respectively. A value < 0 indicates membrane insertion of those residues.

Our canonical MD simulations confirm the peripheral membrane association of the PHD for all protonation states of PIP lipids. In all the mutant simulations, we observed a deeper membrane insertion of the PHD via the E17K residue. In **Supplemental Figure S1 and S2**, we show the residue-wise z-distance for all 24 systems. The z-distance profiles clearly show that membrane insertion happens via Tyr18, Ile19, and Lys20 residues in WT. Lys17 in the mutant system also plays a prominent role. Notably, a longer timescale of the simulation was required to reach the membrane-bound state for WT AKT1-PHD than for mutant systems. For WT configurations, it generally took 750-800 ns to get the bound state, while for the mutant system, membrane association was obtained within 400 ns.

To obtain detailed residue-level information of the PHD involved in membrane binding via PIP lipid, the binding pocket was analyzed for all the bound systems, and the residues within a distance cutoff of 3.5 Å of PIP lipid were considered.^37,38^ These calculations were carried out using the last 50 ns of the simulation to estimate the frequency of occurrence of these residues. Unsurprisingly, a number of these residues belong to Lysines and Arginines, but distinct variation in the PHD residues involved in PIP lipid’s binding pocket was also observed with different protonation states for both WT and E17K systems. Binding pocket residues obtained in the last snapshot of differentially protonated states of PIP_3_ and PIP_2_, along with their corresponding normalized frequency of occurrence calculated over the last 50 ns of trajectory, are shown in **Supplemental Figure S3**. The data reveal the degenerate binding pose across different protonation states.

### AKT1-PHD dissociation free-energies have remarkable variations with changes in protonation states of lipids

From the all-atom simulation trajectories and the z-distance analyses thereafter, where all states showed facile membrane association albeit with different binding geometries, the strength of binding could not be gauged. To quantify the dissociation free energy of each of the states, we performed US-MD simulations to estimate potentials of mean force (PMFs) for the interaction of AKT1 PHD with differently protonated PIP_3_- and PIP_2_-containing lipid bilayers. The starting configuration of this simulation was taken from the bound state of AAMD simulations. We used a collective variable (CV) as the distance between the center of mass of the AKT1-PHD and the membrane midplane. PMF for the eight different protonation states of PIP_3_ lipids are shown in **Figure 5A**. Similarly, PMFs for WT and E17K AKT1-PHD with four different prototation states of PIP_2_ lipids are shown in **Figure 5B**. For the sake of clarity and clear comparisons, we present the ΔG of dissociation for all the systems in **Table 2**.

**Figure 5:**
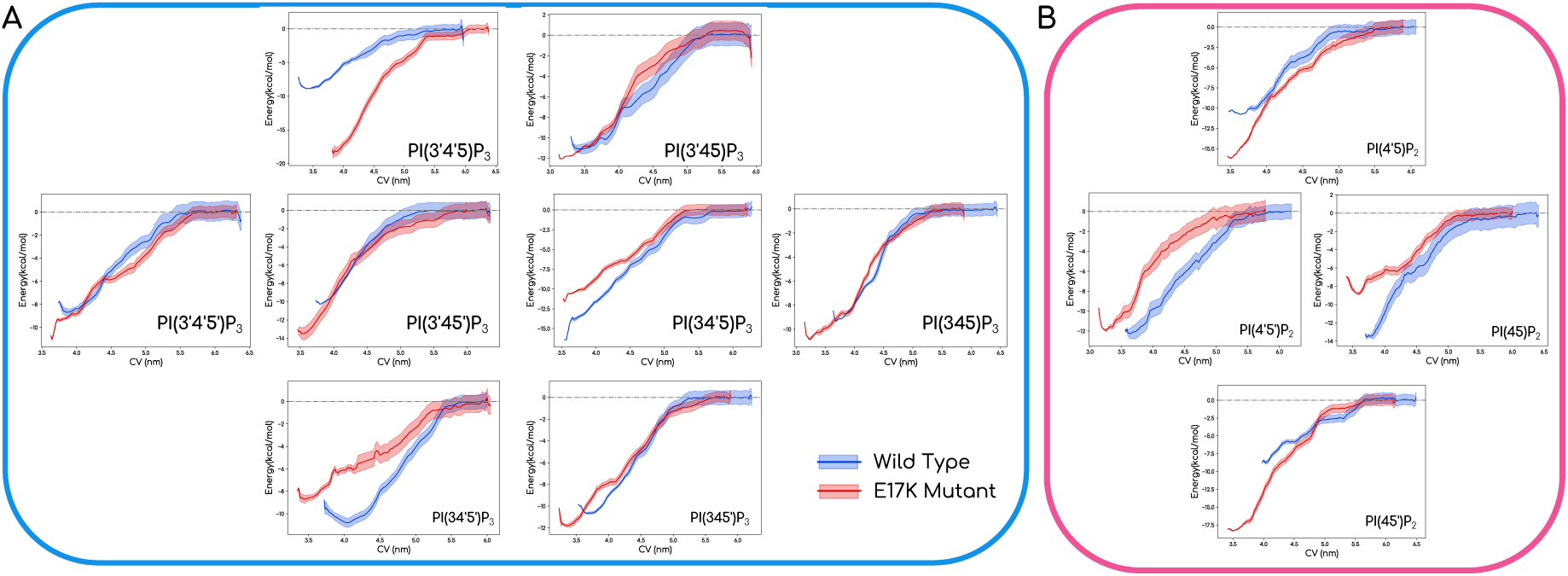
(A) PMF plots of both wildtype and mutant for different protonation states for PIP_3_. (B) PMF plots of both wildtype and mutant for different protonation states for PIP_2_.

**Table 2:** PMF values (ΔG) calculated as a function of the PI3K-AKT1 PH protein-bilayer minimum distance of WT and mutant states for all PIP_3_ and PIP_2_ states. A free energy range of −9.1 to −13.5 kcal/mol was obtained for the WT PHD with a differently protonated PIP_2_ containing bilayer. The values for the corresponding mutant PHD varied within the range of −8.8 to −18.4 kcal/mol.

| Name of PIP<sub>3</sub> lipid | $\Delta G$ Wildtype (kCal) | $\Delta G$ Mutant (kCal) | $\Delta\Delta G_{Mutant-WT}$ (kCal) |
| --- | --- | --- | --- |
| PI(3',4',5')P<sub>3</sub> | -8.846 | -11.12 | -2.274 |
| PI(3,4',5')P<sub>3</sub> | -11.04 | -11.91 | -0.87 |
| PI(3',4,5')P<sub>3</sub> | -10.35 | -13.72 | -3.37 |
| PI(3',4',5)P<sub>3</sub> | -9.264 | -18.46 | -9.196 |
| PI(3,4,5')P<sub>3</sub> | -10.89 | -12.04 | -1.15 |
| PI(3,4',5)P<sub>3</sub> | -14.25 | -11.81 | 2.44 |
| PI(3',4,5)P<sub>3</sub> | -11.16 | -12.46 | -1.3 |
| PI(3,4,5)P<sub>3</sub> | -9.448 | -10.85 | -1.402 |
| PI(4',5')P<sub>2</sub> | -12.29 | -12.04 | 0.25 |
| PI(4,5')P<sub>2</sub> | -9.137 | -18.4 | -9.263 |
| PI(4',5)P<sub>2</sub> | -10.93 | -16.27 | -5.34 |
| PI(4,5)P<sub>2</sub> | -13.47 | -8.834 | 4.636 |

ΔG values exhibit remarkable variations across different protonation states, which is the most revealing part of this study. For example, different protonation states with the same charge (ionization states) showed noticeable changes in ΔG values. Also, WT and E17K showed significant difference in lipid binding affinity for PI(3’,4’,5)P_3_ and PI(4,5’). Also, not all protonation states bind better with mutant systems as shown in the data from (PI(3,4’,5)P_3_ and PI(4,5)P_2_. The ‘z-plots’ (**Figure and Supplemental Figures S1, S2**), which only report the geometric distances of the residues, can often be misleading in terms of the strength of binding. We wanted to understand the molecular interaction origin of the vastly different ΔG values in the 24 different systems. To that end, we calculated the strength and persistence time of the hydrogen bonds that individual residues formed with lipids across the different systems (see **Figure 6**).

**Figure 6:**
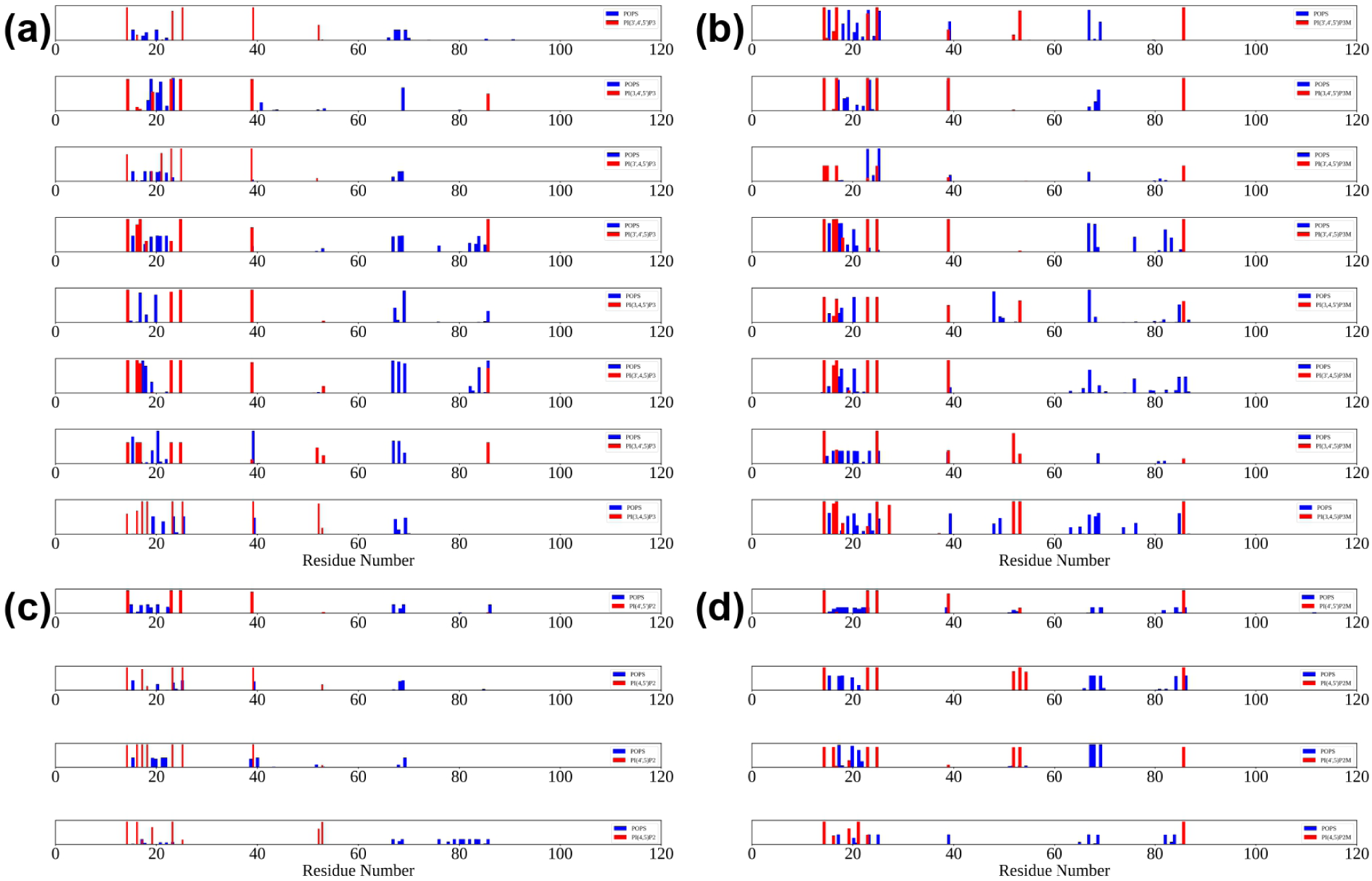
(a,b) Plot showing strength and duration of H-bond formed between POPS and residues of AKT1-PHD shown in blue and PIP_3_ lipid and residues of AKT1-PHD shown in red during the last 50ns of trajectory for both wildtype and mutant states, respectively. (c,d) Plot showing strength and duration of H-bond formed between POPS and residues of AKT1-PHD shown in Blue and PIP_2_ lipid and residues of AKT1-PHD shown in Red during the last 50ns of trajectory for both wildtype and mutant states, respectively.

The H-bond data in **Figure**and ΔG reported in **Table** show consistent correlation. As an example, we compare the profile of PI(3’,4’,5’) and PI(3,4’,5) interactions with WT AKT1-PHD (**Figure 6a)**. Residue-wise H-bond interactions are shown in row 1 and row 6 of **Figure 6a** for PI(3’,4’,5’)_3_ and PI(3,4’,5)_3_, respectively. The differences clearly explain the relatively higher binding strength with the bilayer containing PI(3,4’,5). Similarly, there is a high difference in binding affinity for PI(3’,4’,5) for WT and E17K (see row 4 in (**Figure 6a)** for WT and row 4 (**Figure 6b)** for E17K. There is a higher number of PIP binding residues in E17K, and they are also more persistent. A larger number of residues persistently having an h-bond with POPS lipids in E17K causes a deeper ΔG in PMF readouts. The same is true for PI(4,5’)P_2_ binding data of WT and E17K, where we see a drastic increase in binding affinity for the mutant system in the ΔG calculations. Row 2 in **Figure 6c)** for WT and row 2 in **Figure 6d)** for E17K clearly reveal the origins of increased affinity for E17K mutants. Together, data obtained from US-MD simulations and the H-bond analyses reinforce the highly stereo-specific nature of PIP lipid binding, where repositioning of a single hydrogen/proton (without changing the charge) can have a significant effect on binding strength.

We also wanted to check if the lipid head group orientation changed in the bound state of the protein. We find that the PIPs "necks" out of the member when bound to the AKT1-PHD, which likely facilitates the stereospecific binding as well. Previously, Richard Pastor and co-workers reported that PIP lipids exhibit a distinct inositol ring orientation to the bilayer.^39^ We find that the presence of peripheral proteins has a large effect on ring orientation. To calculate the tilt in the headgroup, we also used the internal vector of the rigid ring structure (see the schematic in **Supplemental Figure S4(a)**. *ϕ* specify the angle (in degrees) made by the C1–C4 vector to the bilayer normal. We wanted to check whether this observed variation is a combined outcome of the difference in protonation state of the lipid and the presence of protein. For this, the tilt angle for the corresponding PIP lipid in the lower leaflet, where the protein was absent, was calculated for comparison. The average values remained almost similar across the protonation states in the unbound PIPs, with the decrease in spread confirming the effect of protein on lipid tilt values, as shown in **Supplemental Figure S4(b)** and **Supplemental Figure S4(c)** for PIP3 and PIP2, respectively. The data from the graphs are reported as a table for convenience in **Supplemental Table S3.**

### Maximum Entropy based integration of NMR-derived populations with US-MD **Δ**G values reconciles the dissociation constant data from biochemical experiments

We use the Maximum Entropy approach to integrate the NMR-derived population data, dissociation constant data from biophysical experiments, and the PMF data from our US-MD simulations. For the sake of clarity, we list out the components(various pieces of information/data used in our approach)

1. The fraction of individual protonation states as a function of pH for both PIP_3_ and PIP_3_ lipids using equation sets 1 and 2, respectively derived using *pK_a_* s from previous NMR-based studies. ^19^
2. The free energy difference between bound and unbound states of AKT-1 PHD calculated from the umbrella sampling simulations in this study 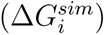.
3. Experimentally determined values of dissociation constants 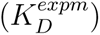 of wild type and mutant AKT1 PHD with lipid membranes^15^ determined at a pH of 7.4. We use these values to estimate the difference in free energies of bound-unbound states using 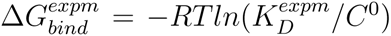.^40^ Here, *R* is the universal gas constant, *T* is the temperature, 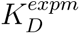 is the experimentally determined dissociation constant of either wild type or mutant AKT1-PHD, *C*^0^ is the standard reference concentration of 1 *M*.

To begin with, we use components 1 and 2 to estimate the simulation-derived free energy of binding 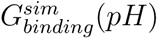 as a function of pH using equation 3, where the summation index *i* in equations 3 and 4 runs over the protonation states of the PIP lipid of interest.

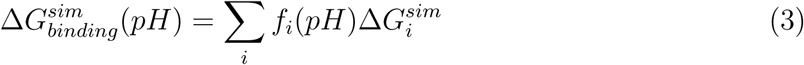

The uncertainty (Δ*G_error_*) on these curves was evaluated using primitive error propagation relations (equation 4), accounting for (i) Uncertainty on fraction of each protonation state (*f_i,error_*(*pH*)) using error evaluated on *pK_a_*s of the above model.^19^ (ii) Uncertainty quantified using bootstrap analysis from pmf profiles 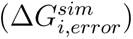 obtained from umbrella sampling simulations.

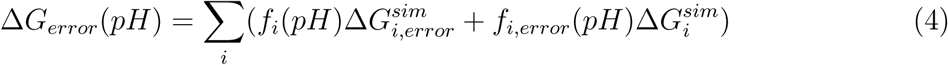

using Equations 3 and 4, we evaluate the free-energy of binding as a function of pH directly and observe the trends as shown in **supplemental Figure S5**. Upon comparing the free energy of binding at pH = 7.4 (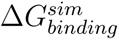 (7.4)) with that from the study listed in component 3 above (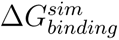 (7.4)), we notice a difference in the estimates. We then probed whether perturbations to the simulation-derived free energies can reconcile the differences observed. We do this carefully using the Bayesian/MaxEnt (BME) reweighting approach based on the principle of maximum entropy,^23^ a method that has recently gained popularity in the context of generating conformational ensembles consistent with experimental data. ^41,42^ We use this approach to reweigh the umbrella sampling derived free energies against the free energy of binding observed at pH = 7.4. The method includes the following constraints

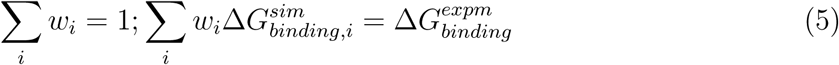

where *w_i_* is the weight/probability of *i*th protonation to the overall experimental 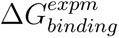 and 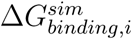 is the free energy difference observed for protonation *i* in simulations. The approach uses the parameter *θ* (parametric temperature). Thus, at higher values of *θ*, entropy dominates. While at lower values of *θ* (*θ →* 0) the method simply fits the simulation data to that of the experimental value (see **Figures 7A and 7B**). A detailed discussion of the parameter *θ* and reweighing the distribution(s) in the light of available data can be found in the literature.^22,23^ The reweighted values for the various systems are shown in **Figure 7C-7E**. We use this framework to reweigh umbrella-sampling derived free energies to calculate the free energies as a function of pH (see **Figures 8A and 8B**).

**Figure 7:**
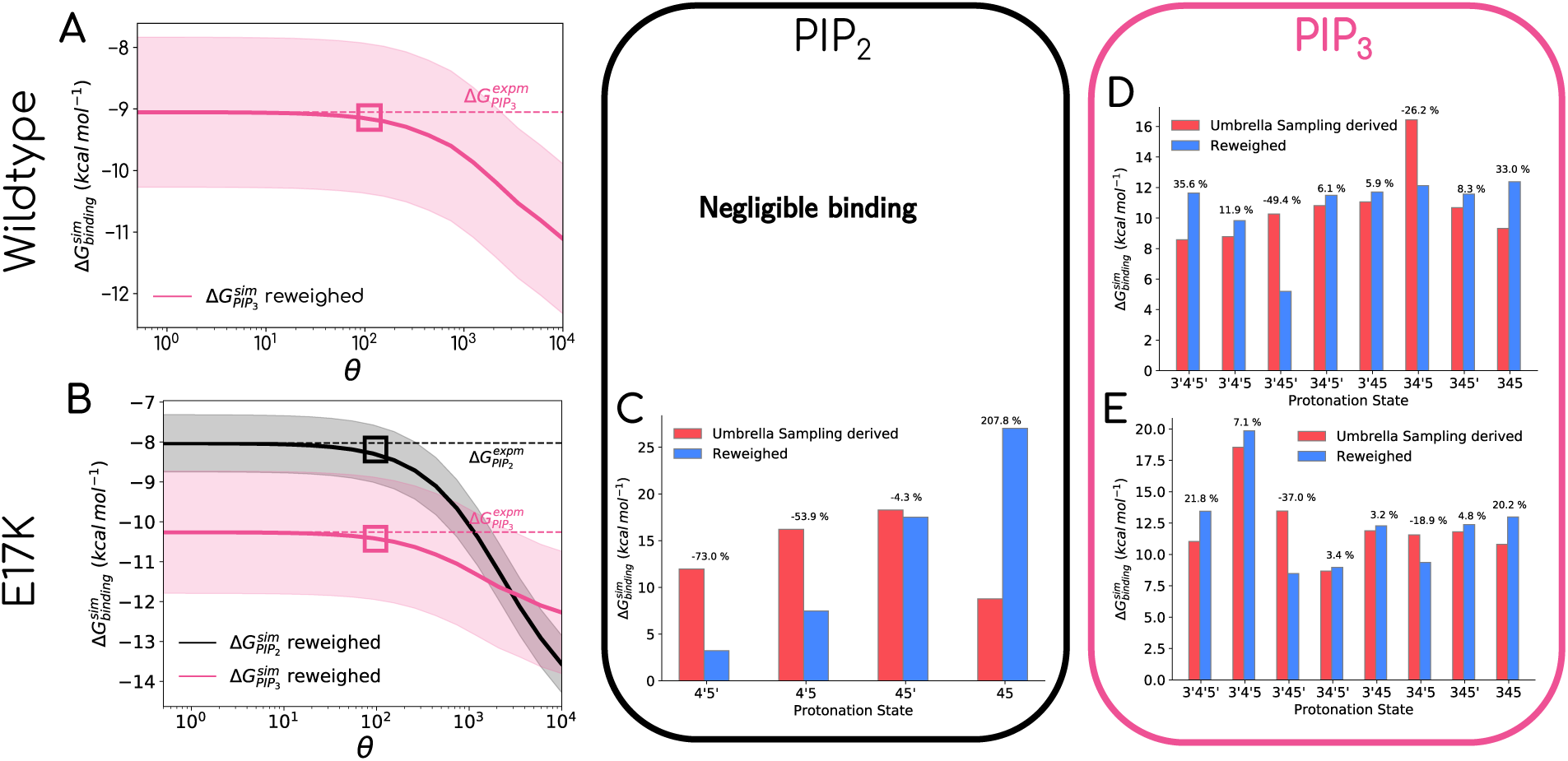
Reweighing Umbrella sampling derived free energies (First row): (A) shows the calculated free energy of binding as a function of *θ* for AKT1-WT with PIP_3_(solid pink line) protonation states. The experimental value observed with this protonation state is marked as a horizontal dotted line for reference. (D) shows the needed change in free energy (shown as blue bars) with respect to those obtained from umbrella sampling (shown as red bars) for PIP_3_ at the *θ* value pointed out in plot (A). (B) Calculated free energy of binding from Umbrella Sampling as a function of *θ* for AKT1-E17 with PIP_3_(solid pink line) protonation states and with PIP_2_(solid black line). The experimental value observed with these lipids is marked as a horizontal dotted lines for PIP_3_(dotted pink line) and PIP_2_(dotted black line). (C) and (E) show the needed change in free energy (shown as blue bars) with respect to those obtained from umbrella sampling (shown as red bars) for PIP_2_ and PIP_3_ at the *θ* value pointed out in plot (B).

**Figure 8:**
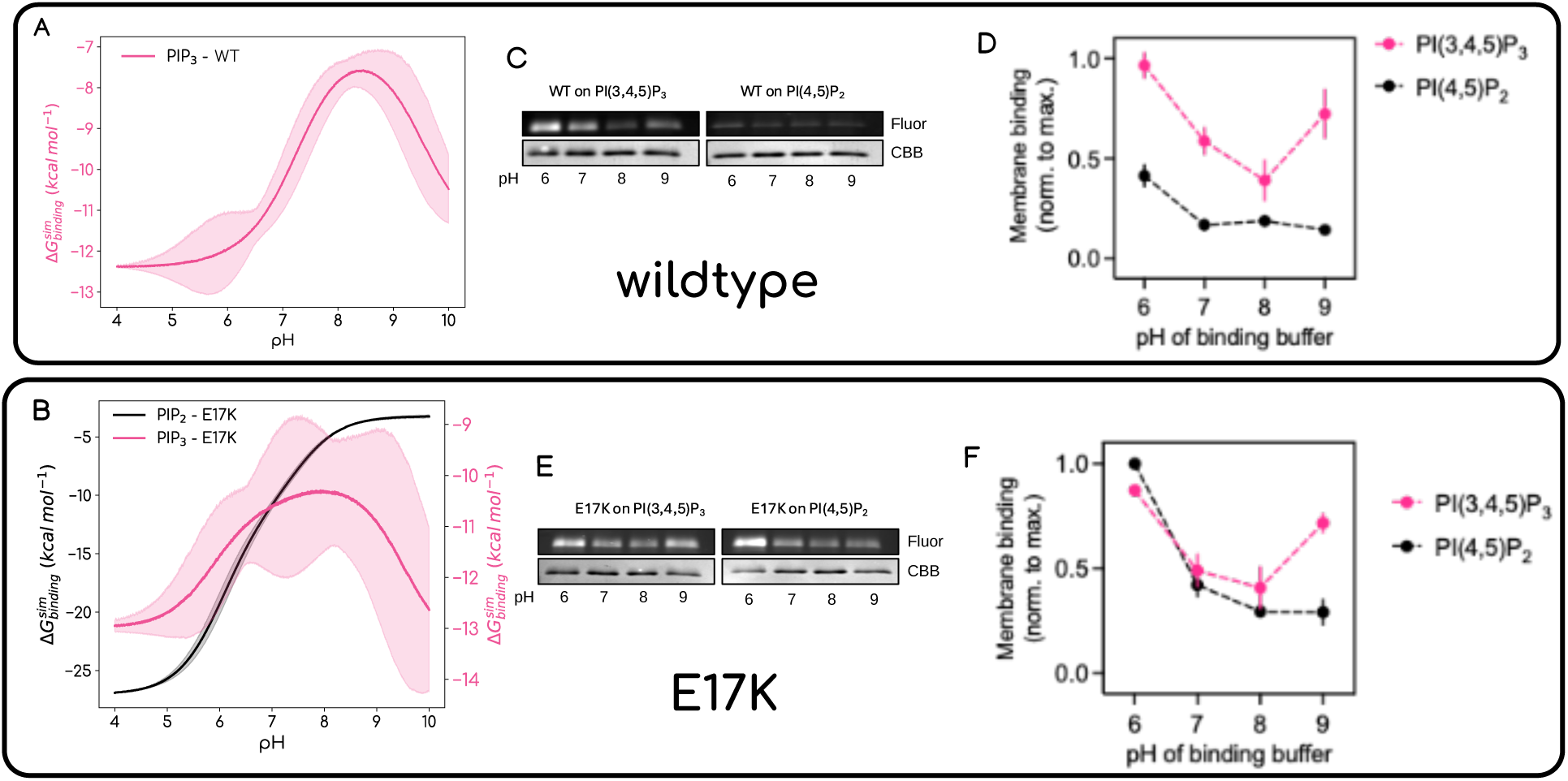
pH differentially modulates the phosphoinositide binding of WT and E17K AKT1 PH domains. Free energy of binding as a function of pH for (A) AKT1-WT and (B) AKT1-E17K mutant with PIP_2_(pink) and PIP_3_(light blue) systems, respectively. (C) Representative PLiMAP assay gels displaying in-gel lipid fluorescence (Fluor) and Coomassie Brilliant Blue (CBB) total protein staining of the wild-type (WT) AKT1 PH domain bound to vesicles containing either PI(3,4,5)P_3_ or PI(4,5)P_2_ across a pH range of 6.0–9.0. (D) Quantification of WT AKT1 PH domain membrane binding to PI(3,4,5)P_3_ (magenta) and PI(4,5)P_2_ (black) vesicles as a function of buffer pH. (E) Representative PLiMAP assay gels displaying in-gel lipid fluorescence (Fluor) and CBB total protein staining of the oncogenic mutant E17K AKT1 PH domain bound to PI(3,4,5)P_3_ or PI(4,5)P_2_ vesicles across a pH range of 6.0–9.0. (F) Quantification of E17K AKT1 PH domain membrane binding to PI(3,4,5)P_3_ (magenta) and PI(4,5)P_2_ (black) vesicles as a function of buffer pH. Data in (D) and (F) represent the mean ± SD of three independent experiments. Fluorescence was normalized to the sample showing maximum binding, which was with WT on PI(3,4,5)P_3_ at pH 6.0 and the E17K mutant on PI(4,5)P_2_ at pH 6.0.

### PliMAP experimental assays confirm that AKT1 PH domain binding to distinct phosphoinositide species is differentially sensitive to pH

**Figure 8(A)** shows the predicted free energy of binding as a function of pH for AKT1-WT for PIP_3_ containing membrane. The data shows that the binding follows a bell shaped behavior as a function of pH with poorest binding in middle and lower/higher pH showing better binding. **Figure 8(B)**) shows the predicted free energy of binding as a function of pH for AKT1-E17K mutant systems for PIP_3_ containing membrane (Pink) and PIP_2_ containing membrane (Black). E17K PH Domain PIP_3_ binding again shows a bell-shaped binding behavior while PIP_2_ lipid binding follows a sigmoidal profile with best binding at low pH and poorer binding at higher pH values. To experimentally confirm our simulations-based predictions, we utilized our Proximity-based Labeling of Membrane Associated Proteins (PLiMAP) assay^34,35^ to analyze AKT1 PH domain binding to vesicles with PIP_3_ and vesicles with PIP_2_ lipids for both WT and E17K PH domains. Notably, the wild-type (WT) AKT1 PH domain selectively binds the rare signaling lipid PI(3,4,5)P_3_, which is generated at the plasma membrane upon PI3K pathway activation. Our first system was PLiMAP assay with WT AKT1 PH domain binding to vesicles composed of 1 mol% PI(3,4,5)P_3_, 98 mol% phosphatidyl-choline (PC), embedded with 1 mol% of a bifunctional, fluorescent, crosslinking lipid probe. In this system, membrane binding positions the protein adjacent to the probe, leading to covalent photo-crosslinking upon UV exposure. Samples are subsequently resolved via SDS-PAGE, and membrane binding is quantified by measuring the lipid-associated fluorescence of the protein band.

Because fluctuations in pH alter the protonation states of phosphoinositides—potentially modulating protein-lipid interactions as show from our computational work — we evaluated binding across a pH range of 6.0 to 9.0. Notably, the AKT1 PH domain exhibited a distinct, pH-dependent binding profile toward PI(3,4,5)P_3_-containing vesicles (**Figure 8(C,D)**). This interaction followed a bell-shaped curve - binding peaked at pH 6.0, decreased at pH 7.0 and 8.0, and rose again at pH 9.0. These data demonstrate that pH variations directly influence the interaction between WT AKT1 and PI(3,4,5)P3. In contrast, control experiments monitoring binding to PI(4,5)P_2_ revealed that while binding was also maximal at pH 6.0, it decreased at pH 7.0 and plateaued through pH 8.0 and 9.0. Taken together, these results indicate that pH differentially regulates AKT1 PH domain interactions across distinct phosphoinositide species.

The oncogenic E17K mutation within the AKT1 PH domain has been reported to enhance binding to the highly abundant lipid PI(4,5)P_2_. Consistently, our PLiMAP assay confirmed that the E17K mutant exhibits better binding to PI(4,5)P_2_compared to the WT protein (**Figures 8(E,F)**). Notably, this mutant–PI(4,5)P_2_ interaction also showed a decline with an increase in pH. In stark contrast, the binding of the E17K mutant to PI(3,4,5)P_3_ displayed a pronounced pH dependence mirroring that of the WT protein, a characteristic notably absent in its interaction with PI(4,5)P_2_. Together, these results demonstrate that environmental pH exerts a significant, lipid-specific influence on both WT and mutant AKT1 PH domains, selectively modulating their interactions with PI(3,4,5)P_3_ but not with PI(4,5)P_2_.

### Conclusions

In summary, this study demonstrates that the membrane association dynamics and dissociation free energies of the AKT1 PH domain are profoundly regulated by the protonation states of phosphoinositide headgroups. By integrating QM-derived partial charges, microsecond-scale all-atom molecular dynamics, and umbrella sampling free-energy calculations with ssNMR-derived lipid populations via Bayesian Maximum Entropy reweighting, we show that subtle variations in the degree and positional placement of protons, rather than simple net charge changes, dramatically alter headgroup orientations, interaction networks, and binding thermodynamics. This integrative framework reconciles long-standing discrepancies across experimental binding assays, providing a clear molecular rationale for the strict PIP_3_ specificity of WT-AKT1 over PIP_2_ under physiological conditions.

Furthermore, our model provides a structural mechanism for the oncogenic hyperactivation caused by the pathological E17K sentry mutation. The single charge reversal at position 17 establishes robust electrostatic interactions that bypass the selective protonation barriers present in wild-type binding, enabling high-affinity interactions with the abundant *PIP*_2_ pool. Direct experimental PLIMAP assays confirmed these predictions, showing a distinct bell-shaped, pH-dependent binding profile for PIP_3_-containing membranes in both WT and E17K AKT1-PHD, whereas the gain-of-function E17K–PIP_2_ interaction decays exponetially as a function of increading pH. These experimental readouts validate that lipid protonation dynamics act as functional regulators of protein-membrane association rather than static bystanders.

In broader physiological terms, these findings highlight local pH fluctuations and lipid microenvironments as fundamental, unappreciated regulators of the PI3K/AKT/mTOR signaling axis. Because different subcellular organelles and pathological microenvironments maintain distinct ionic and pH profiles, the dynamic protonation state of second-messenger lipids offers a spatial mechanism to fine-tune signal transduction. Accounting for these dynamic protonation states and headgroup charge redistributions will be essential for future quantitative modeling of peripheral membrane protein recruitment and the rational design of therapeutics targeting lipid-binding domains.

## Author contributions

Anand Srivastava (AS) conceived the idea and designed the experiments in consultation with Kirtika Jha (KJ) and Krishnakanth Baratam (KB). Jayashree Nagesh performed the quantum calculations for charge and geometry assignment to PIPx lipid headgroups. KJ set up all the MD-based simulations and performed the analyses. KB rendered the final figures and performed integration of simulation and experimental data using the max-entropy approach. Megahdeepa Sarkar performed the pH-dependent binding assay experiments under the supervision of Thomas Pucadyil. Kirtika Jha and Anand Srivastava wrote the article with the help of other authors.

## Acknowledgements

AS acknowledges the financial support from the Indian Institute of Science (IISc) and the high-performance computing facility "Beagle" that was set up from grants by the erstwhile IISc-DBT partnership programme. AS acknowledges the FIST program sponsored by the Department of Science and Technology, India that supports the MBU infrastructure. AS thanks the DBT National Network Project (NNP) grant (BT/PR40323/BTIS/137/78/2023), the Matrics grants (MTR/2023/001040) and the ARG grant (ANRF/ARG/2025/009141/LS)from ANRF, India. AS and TJP would also like to thank the Teams Science Grant from the DBT-Wellcome Trust India Alliance (Grant number: IA/TSG/21/1/600245). TJP thanks the Howard Hughes Medical Institute for an International Research Scholar’s Grant (Grant No. 55008746), the Anusandhan National Research Foundation for a SUPRA Grant (Grant No. SPR12021100014). KB thanks ANRF for the National Postdoctoral Fellowship Grant (ANRF/PDF/2025/006111). MS thanks the CSIR for graduate student PhD fellowship.

## Supporting Information

**Table S1:** Charges obtained from *ab initio* QM calculations of headgroup atoms of various protonation states of PIP_3_ lipid.

| Atom Type | Partial Charge on each atom type |  |  |  |
| --- | --- | --- | --- | --- |
|  | PI(3,4,5)P<sub>3</sub> | PI(3',4,5)P<sub>3</sub> | PI(3,4',5)P<sub>3</sub> | PI(3,4,5')P<sub>3</sub> |
| C11 | 0.298 | 0.269 | 0.284 | 0.56 |
| C12 | 0.137 | 0.197 | 0.522 | 0.425 |
| C13 | -0.227 | -0.596 | -1.248 | -1.208 |
| C14 | 1.098 | 0.815 | 1.334 | 1.091 |
| C15 | -1.251 | -0.229 | -0.262 | -0.096 |
| C16 | 0.512 | 0.175 | 0.229 | -0.073 |
| H1 | 0.086 | 0.084 | 0.074 | 0.028 |
| H2 | 0.12 | 0.227 | 0.163 | 0.177 |
| H3 | 0.172 | 0.248 | 0.349 | 0.351 |
| H4 | -0.011 | 0.031 | -0.065 | -0.041 |
| H5 | 0.262 | 0.193 | 0.156 | 0.143 |
| H6 | 0.371 | 0.123 | 0.083 | 0.136 |
| O2 | -0.758 | -0.721 | -0.785 | -0.786 |
| HO2 | 0.518 | 0.456 | 0.483 | 0.508 |
| O3 | -0.723 | -0.508 | -0.307 | -0.288 |
| P3 | 1.326 | 1.523 | 1.356 | 1.319 |
| OP32 | -0.913 | -1.06 | -0.671 | -0.646 |
| OP33 | -0.652 | -1.017 | -0.928 | -0.915 |
| OP34 | -0.909 | -1.029 | -0.919 | -0.909 |
| HP32 | 0.462 |  | 0.462 | 0.459 |
| O4 | -0.689 | -0.652 | -0.891 | -0.696 |
| P4 | 1.481 | 1.519 | 1.787 | 1.503 |
| OP44 | -0.972 | -0.719 | -1.092 | -0.981 |
| OP43 | -0.933 | -0.953 | -1.063 | -0.947 |
| OP42 | -0.732 | -0.99 | -1.114 | -0.74 |
| HP42 | 0.481 | 0.48 |  | 0.478 |
| O5 | -0.284 | -0.729 | -0.766 | -0.736 |
| P5 | 1.684 | 1.668 | 1.695 | 1.791 |
| OP52 | -0.748 | -0.75 | -0.742 | -1.084 |
| OP53 | -0.968 | -0.964 | -0.972 | -1.09 |
| OP54 | -0.962 | -0.957 | -0.972 | -1.074 |
| HP52 | 0.494 | 0.496 | 0.477 |  |
| O6 | -0.656 | -0.769 | -0.796 | -0.778 |
| HO6 | 0.472 | 0.515 | 0.54 | 0.567 |
| O12 | -0.422 | -0.403 | -0.435 | -0.456 |
| P | 0.92 | 0.929 | 0.933 | 0.92 |
| O13 | -0.616 | -0.611 | -0.613 | -0.632 |
| C1 | 0.464 | 0.463 | 0.471 | 0.476 |
| HB | -0.012 | -0.008 | -0.015 | -0.018 |
| HA | -0.002 | -0.006 | -0.004 | -0.006 |
| C2 | -0.374 | -0.354 | -0.37 | -0.368 |
|  | PI(3',4',5)P<sub>3</sub> | PI(3',4,5')P<sub>3</sub> | PI(3,4',5')P<sub>3</sub> | PI(3',4',5')P<sub>3</sub> |
| C11 | 0.349 | 0.628 | 0.563 | 0.617 |
| C12 | 0.21 | -0.058 | 0.417 | 0.038 |
| C13 | -0.781 | 0.011 | -1.165 | -0.061 |
| C14 | 1.292 | 1.082 | 1.444 | 1.258 |
| C15 | -0.149 | -0.717 | -0.297 | -0.74 |
| C16 | 0.238 | 0.104 | 0.034 | 0.12 |
| H1 | 0.048 | -0.002 | 0.02 | -0.008 |
| H2 | 0.231 | 0.116 | 0.173 | 0.083 |
| H3 | 0.26 | 0.082 | 0.274 | 0.079 |
| H4 | -0.103 | -0.087 | -0.101 | -0.121 |
| H5 | 0.096 | 0.257 | 0.176 | 0.247 |
| H6 | 0.066 | 0.244 | 0.105 | 0.238 |
| O2 | -0.739 | -0.8 | -0.798 | -0.834 |
| HO2 | 0.456 | 0.573 | 0.509 | 0.594 |
| O3 | -0.47 | -0.76 | -0.462 | -0.793 |
| P3 | 1.469 | 1.443 | 1.569 | 1.462 |
| OP32 | -1.035 | -1.002 | -0.745 | -1.014 |
| OP33 | -1.013 | -1.022 | -0.966 | -1.034 |
| OP34 | -1.033 | -1.025 | -0.941 | -1.033 |
| HP32 |  |  | 0.464 |  |
| O4 | -0.897 | -0.724 | -0.915 | -0.888 |
| P4 | 1.734 | 1.465 | 1.806 | 1.742 |
| OP44 | -1.068 | -0.958 | -1.105 | -1.075 |
| OP43 | -1.045 | -0.934 | -1.097 | -1.054 |
| OP42 | -1.126 | -0.737 | -1.063 | -1.136 |
| HP42 |  | 0.465 |  |  |
| O5 | -0.788 | -0.463 | -0.774 | -0.471 |
| P5 | 1.713 | 1.804 | 1.835 | 1.835 |
| OP52 | -0.752 | -1.088 | -1.099 | -1.102 |
| OP53 | -0.98 | -1.096 | -1.092 | -1.103 |
| OP54 | -0.982 | -1.083 | -1.096 | -1.098 |
| HP52 | 0.481 |  |  |  |
| O6 | -0.814 | -0.739 | -0.915 | -0.745 |
| HO6 | 0.544 | 0.48 | 0.603 | 0.468 |
| O12 | -0.439 | -0.46 | -0.473 | -0.476 |
| P | 0.947 | 0.934 | 0.925 | 0.947 |
| O13 | -0.618 | -0.636 | -0.449 | -0.635 |
| C1 | 0.48 | 0.483 | 0.472 | 0.493 |
| HB | -0.015 | -0.017 | -0.006 | -0.02 |
| HA | -0.01 | -0.011 | -0.015 | -0.014 |
| C2 | -0.353 | -0.351 | -0.365 | -0.348 |

**Table S2:** Charges obtained from *ab initio* QM calculations of headgroup atoms of various protonation states of PIP_2_ lipid.

| Atom type | Partial Charge on each atom type |  |  |  |
| --- | --- | --- | --- | --- |
|  | PI(4,5')P<sub>2</sub> | PI(4',5)P<sub>2</sub> | PI(4',5')P<sub>2</sub> | PI(4,5)P<sub>2</sub> |
| C12 | -0.72 | 0.633 | 0.564 | 0.74 |
| H2 | 0.273 | 0.107 | 0.128 | 0.099 |
| O2 | -0.564 | -0.746 | -0.736 | -0.752 |
| HO2 | 0.452 | 0.425 | 0.42 | 0.449 |
| C13 | 0.645 | 0.035 | -0.043 | -0.007 |
| H3 | 0.028 | 0.144 | 0.133 | 0.153 |
| O3 | -0.765 | -0.888 | -0.893 | -0.861 |
| HO3 | 0.473 | 0.574 | 0.566 | 0.592 |
| C14 | 0.205 | 0.185 | 0.579 | -0.15 |
| H4 | 0.086 | 0.068 | 0.01 | 0.191 |
| O4 | -0.641 | -0.689 | -0.69 | -0.453 |
| P4 | 1.439 | 1.603 | 1.424 | 1.426 |
| OP43 | -0.915 | -1.003 | -0.989 | -0.916 |
| OP44 | -0.909 | -1.043 | -1.052 | -0.931 |
| OP42 | -0.736 | -1.036 | -0.962 | -0.729 |
| HP42 | 0.441 |  |  | 0.501 |
| C15 | 0.079 | 0.049 | -0.106 | -0.084 |
| H5 | 0.051 | 0.128 | 0.153 | 0.17 |
| O5 | -0.577 | -0.637 | -0.646 | -0.605 |
| P5 | 1.537 | 1.485 | 1.54 | 1.472 |
| OP54 | -1.046 | -0.958 | -1.016 | -0.747 |
| OP52 | -1.035 | -0.937 | -1.07 | -0.946 |
| OP53 | -0.997 | -0.762 | -1.059 | -0.926 |
| HP52 |  | 0.481 |  | 0.478 |
| C16 | 0.154 | 0.9 | 0.69 | 1.04 |
| H6 | 0.124 | -0.001 | 0.08 | -0.006 |
| O6 | -0.686 | -0.835 | -0.804 | -0.844 |
| HO6 | 0.416 | 0.439 | 0.438 | 0.44 |
| C11 | 0.314 | -0.987 | -0.867 | -1.074 |
| H1 | 0.147 | 0.313 | 0.299 | 0.343 |
| O12 | -0.492 | -0.282 | -0.307 | -0.294 |
| P | 1.272 | 1.213 | 1.197 | 1.189 |
| O11 | -0.512 | -0.528 | -0.593 | -0.519 |
| O13 | -0.831 | -0.741 | -0.521 | -0.751 |
| O14 | -0.596 | -0.574 | -0.76 | -0.584 |
| C1 | 0.396 | 0.382 | 0.402 | 0.378 |
| HB | 0.031 | 0.033 | 0.028 | 0.037 |
| HA | 0.02 | 0.036 | 0.025 | 0.035 |
| C2 | -0.444 | -0.45 | -0.468 | -0.484 |

**Table S3:** Tilt angle (*ϕ*) of PIP lipid in both Wildtype and mutant bound AKT1 PHD states as well as free state for all protonation states of PIP_3_ and PIP_2_.

| Name of PIP lipid | $\phi$ *Wildtype* | $\phi$ *Mutant* | $\phi$ *Free* |
| --- | --- | --- | --- |
| PI(3',4',5')P<sub>3</sub> | 135.97 ± 5.63 | 162.99 ± 5.07 | 24.98 ± 11.43 |
| PI(3,4',5')P<sub>3</sub> | 133.13 ± 6.18 | 110.42 ± 5.73 | 25.31 ± 10.23 |
| PI(3',4,5')P<sub>3</sub> | 102.07 ± 5.77 | 139.90 ± 4.945 | 34.95 ± 11.85 |
| PI(3',4',5)P<sub>3</sub> | 137.19 ± 5.21 | 128.93 ± 4.84 | 34.46 ± 13.97 |
| PI(3,4,5')P<sub>3</sub> | 143.43 ± 4.186 | 132.39 ± 7.73 | 27.63 ± 14.96 |
| PI(3,4',5)P<sub>3</sub> | 110.68 ± 5.92 | 147.62 ± 7.66 | 44.08 ± 14.37 |
| PI(3',4,5)P<sub>3</sub> | 145.31 ± 6.42 | 111.98 ± 6.08 | 27.10 ± 12.61 |
| PI(3,4,5)P<sub>3</sub> | 137.88 ± 6.03 | 144.57 ± 6.20 | 57.78 ± 16.05 |
| PI(4',5')P<sub>2</sub> | 83.52 ± 5.70 | 120.87 ± 5.88 | 50.48 ± 15.86 |
| PI(4,5')P<sub>2</sub> | 108.20 ± 4.97 | 130.91 ± 5.91 | 36.88 ± 13.64 |
| PI(4',5)P<sub>2</sub> | 120.75 ± 6.12 | 101.23 ± 6.49 | 42.27 ± 17.46 |
| PI(4,5)P<sub>2</sub> | 76.83 ± 5.37 | 118.3 ± 5.21 | 50.56 ± 14.175 |

**Figure S1:**
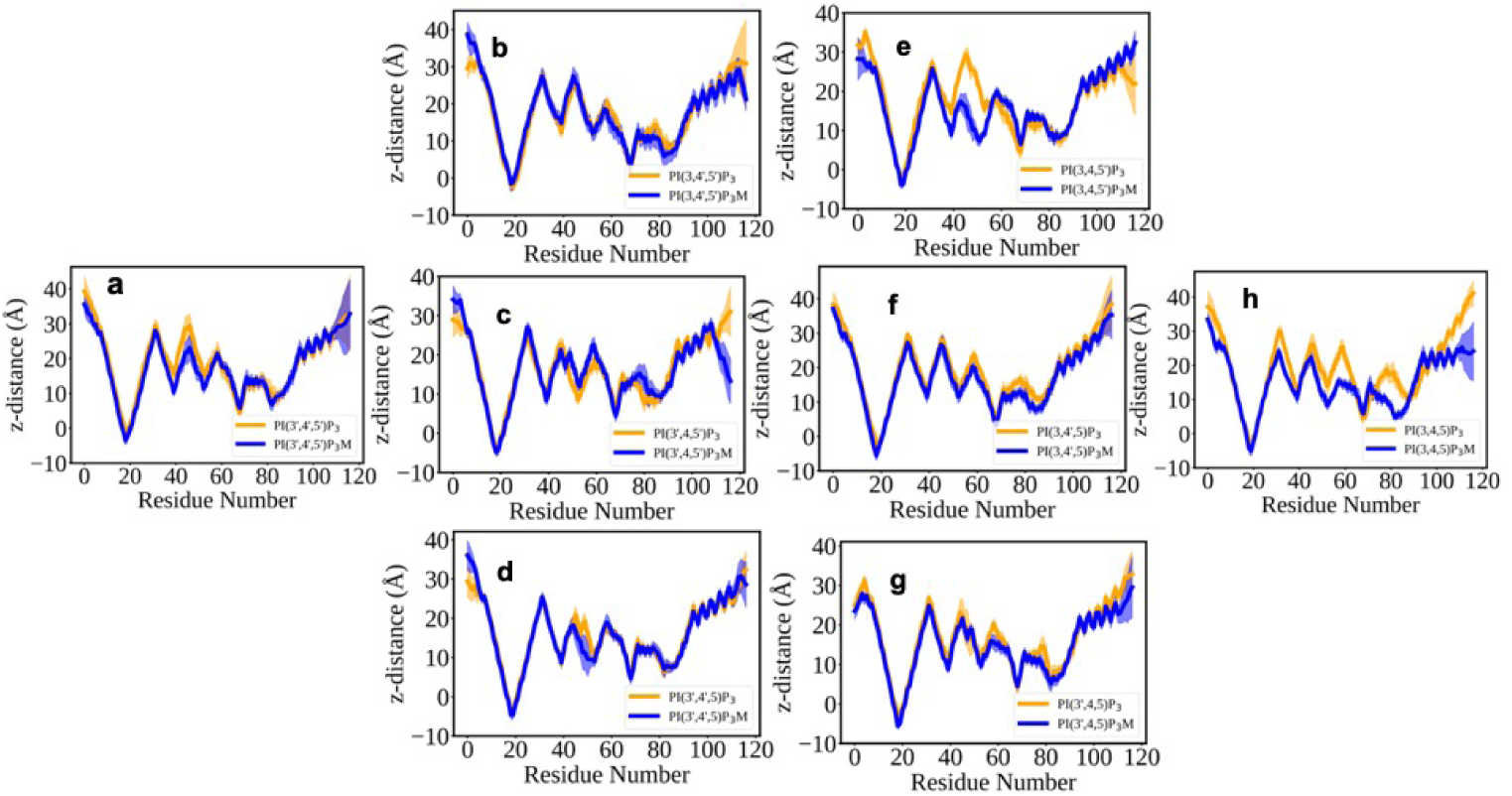
Interaction of AKT1-PHD with differently protonated PIP_3_ containing lipid bilayer. Ensemble-averaged C profile of the bound AKT1-PHD, calculated during last the 50 ns of atomistic simulations. The blue color corresponds to Wildtype and orange to the mutant, respectively. a) PI(3’,4’,5’)P_3_ (No protonation) b) PI(3,4’,5’)P_3_ c) PI(3’,4,5’)P_3_ d) PI(3’,4’,5)P_3_ (Single protonation) e) PI(3,4,5’)P_3_ f) PI(3,4’,5)P_3_ g) PI(3,4’,5’)P_3_ (Double protonation) h) PI(3,4,5)P_3_ (Triple protonation)

**Figure S2:**
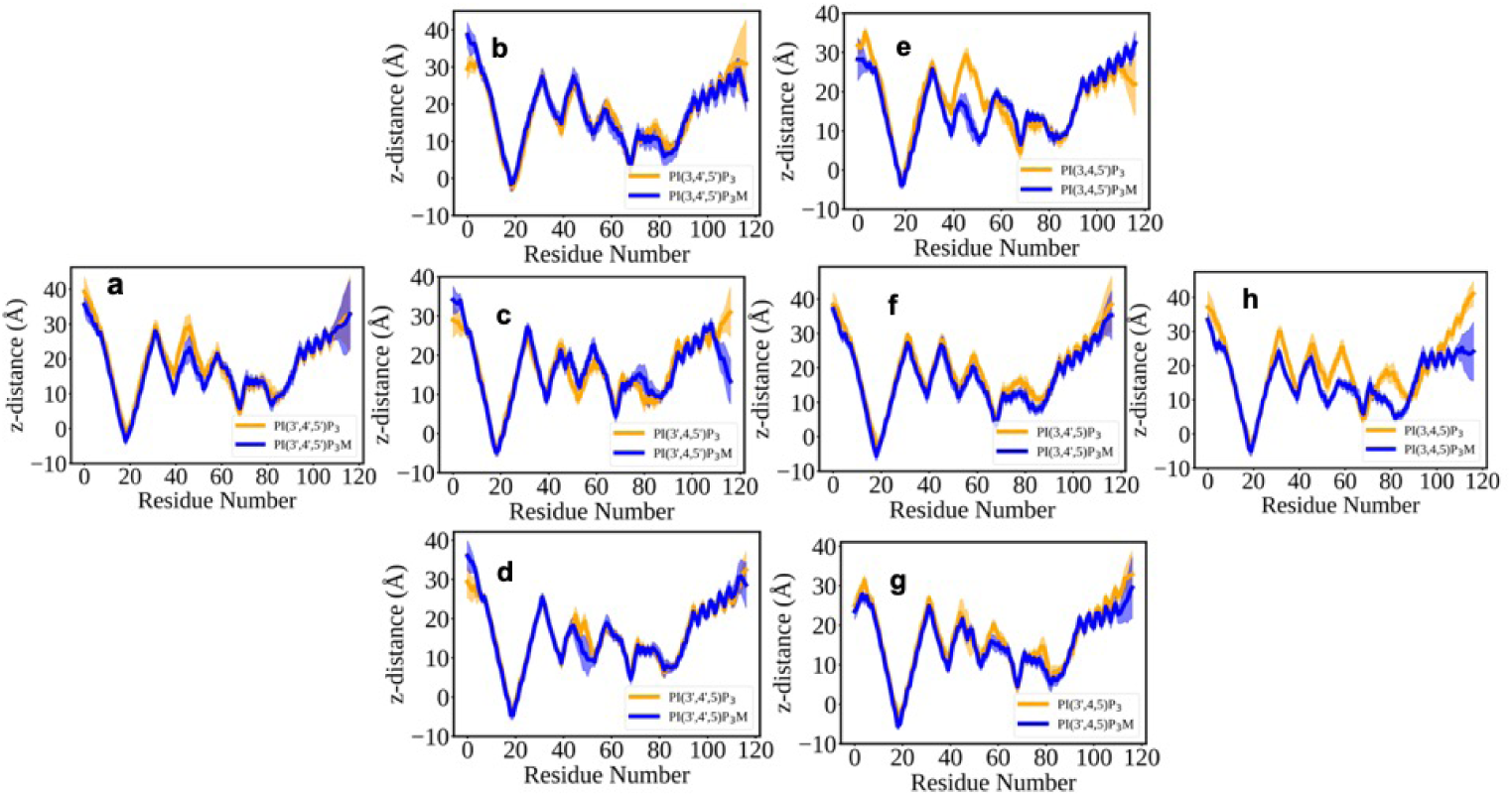
Interaction of AKT1-PHD with differently protonated PIP_3_ containing lipid bilayer. Ensemble-averaged C profile of the bound AKT1-PHD, calculated during last the 50 ns of atomistic simulations. The blue color corresponds to Wildtype and orange to the mutant, respectively. a) PI(4’,5’)P_2_(No protonation) b) PI(4,5’)P_2_ c) PI(4’,5)P_2_ (Single protonation) d) PI(4,5)P_2_ (Double protonation)

**Figure S3:**
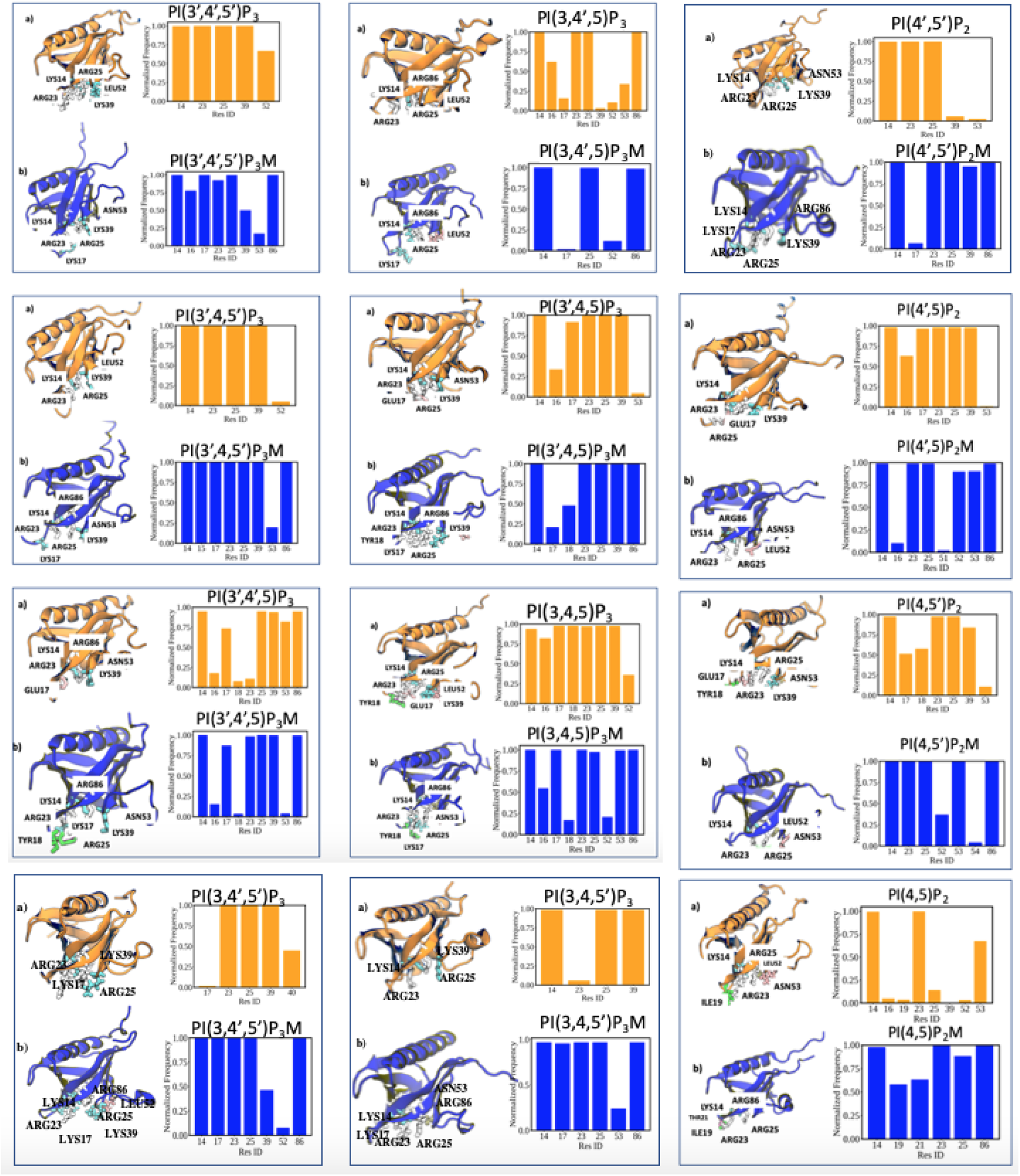
Binding pocket residues obtained in the last snapshot of differently protonated states of PIP_3_ (Left and middle column) and PIP_2_ (Right column) with their corresponding normalized frequency of occurrence calculated over last 50 ns of trajectory. (a) Wildtype PHD (shown in orange) (b) Mutant PHD (shown in blue). AKT1 PHD is shown in cartoon representation.

**Figure S4:**
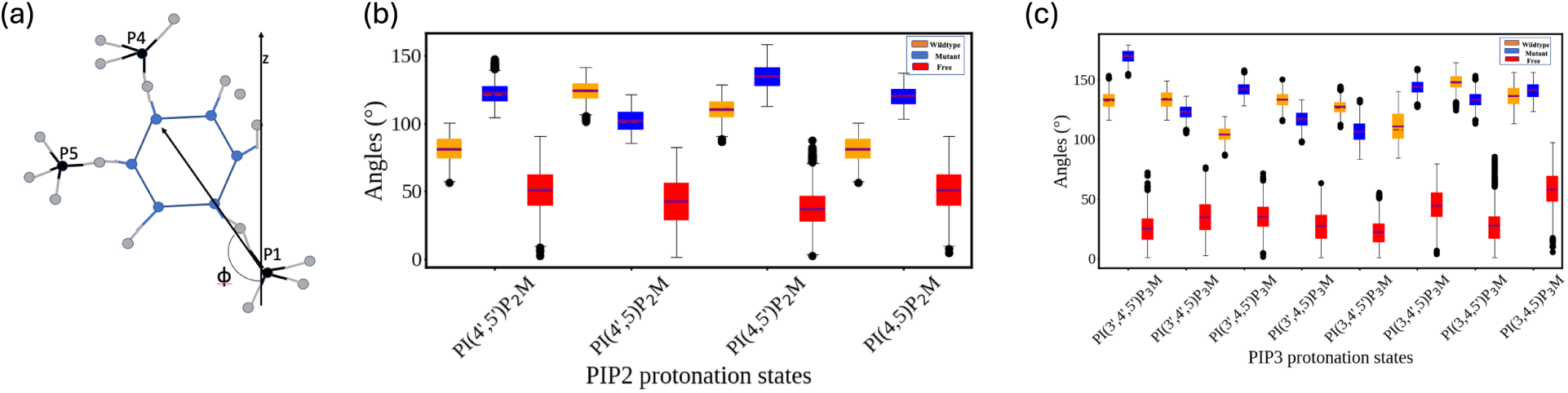
Effect of presence of protein on tilt angle (*ϕ*). (a) Schematic of the tilt angle (*ϕ*) calculation of PIP lipid. (b) The tilt angle of PIP_3_ lipid **Wildtype** PHD bound state shown in orange, **Mutant** PHD bound state shown in blue and **Free** state devoid of protein (lower leaflet of bilayer) shown in red (c) The tilt angle of PIP_2_ lipid **Wildtype** PHD bound state shown in orange, **Mutant** PHD bound state shown in blue and **Free** state devoid of protein (lower leaflet) shown in red

**Figure S5:**
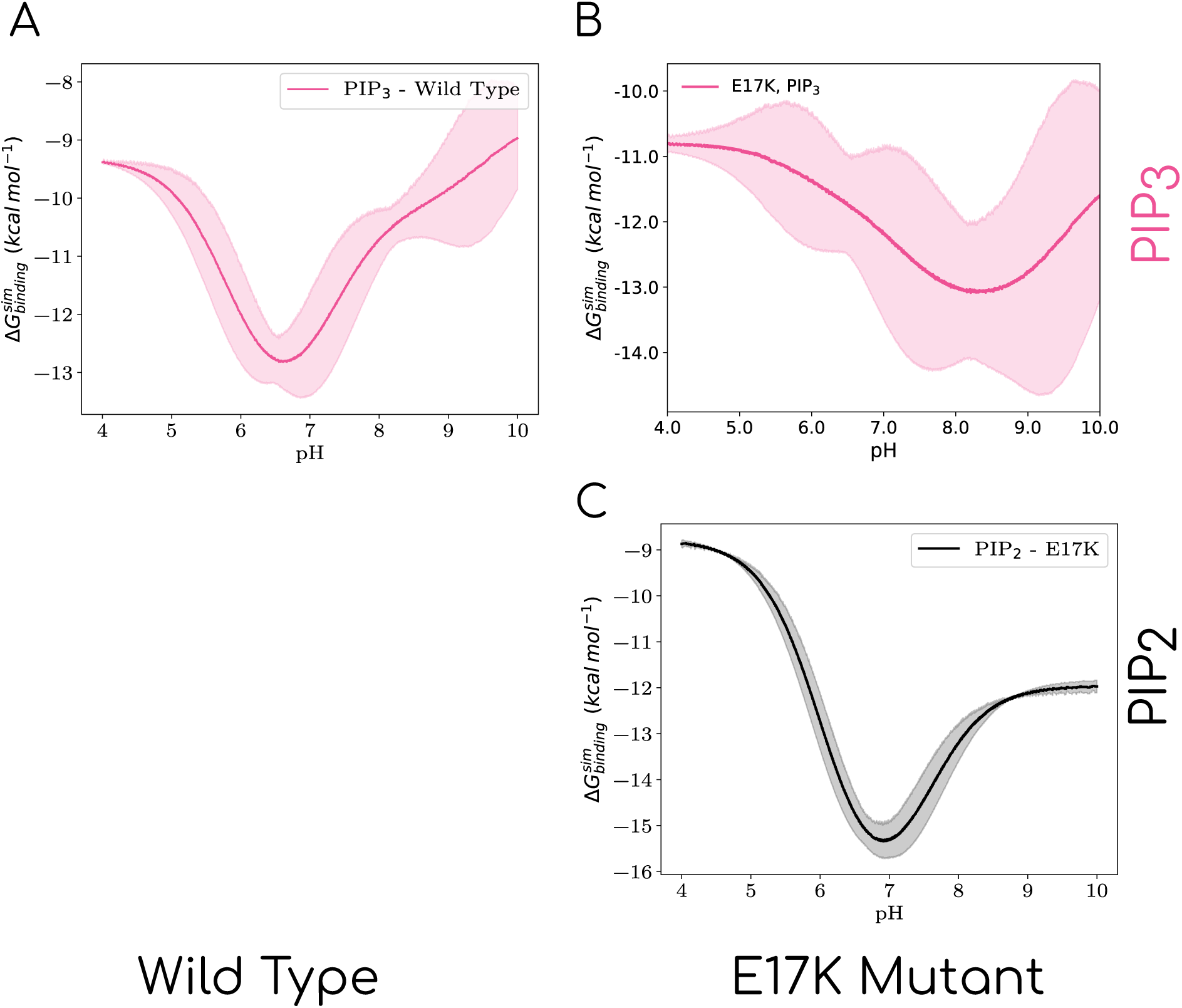
(A) Calculated free energies of binding as a function of pH for wild type with PIP_3_ lipid. (B) Calculated free energies of binding as a function of pH for E17K Mutant with PIP_3_ lipid and (C) PIP_2_ lipid.

## References

(1) Liu, P.; Cheng, H.; Roberts, T. M.; Zhao, J. J. Targeting the phosphoinositide 3-kinase pathway in cancer. Nature reviews Drug discovery 2009, 8, 627–644.

(2) Engelman, J. A. Targeting PI3K signalling in cancer: opportunities, challenges and limitations. Nature Reviews Cancer 2009, 9, 550–562.

(3) Goldar, S.; Khaniani, M. S.; Derakhshan, S. M.; Baradaran, B. Molecular mechanisms of apoptosis and roles in cancer development and treatment. Asian Pac J Cancer Prev 2015, 16, 2129–2144.

(4) Fruman, D. A.; Chiu, H.; Hopkins, B. D.; Bagrodia, S.; Cantley, L. C.; Abraham, R. T. The PI3K pathway in human disease. Cell 2017, 170, 605–635.

(5) Brazil, D. P.; Hemmings, B. A. Ten years of protein kinase B signalling: a hard Akt to follow. Trends in biochemical sciences 2001, 26, 657–664.

(6) Adams, J. R.; Schachter, N. F.; Liu, J. C.; Zacksenhaus, E.; Egan, S. E. Elevated PI3K signaling drives multiple breast cancer subtypes. Oncotarget 2011, 2, 435.

(7) Downward, J. Mechanisms and consequences of activation of protein kinase B/Akt. Current opinion in cell biology 1998, 10, 262–267.

(8) Manning, B. D.; Toker, A. AKT/PKB signaling: navigating the network. Cell 2017, 169, 381–405.

(9) Frech, M.; Andjelkovic, M.; Ingley, E.; Reddy, K. K.; Falck, J. R.; Hemmings, B. A. High affinity binding of inositol phosphates and phosphoinositides to the pleckstrin homology domain of RAC/protein kinase B and their influence on kinase activity. Journal of Biological Chemistry 1997, 272, 8474–8481.

(10) James, S. R.; Peter Downes, C.; Gigg, R.; Grove, S. J.; Holmes, A. B.; Alessi, D. R. Specific binding of the Akt-1 protein kinase to phosphatidylinositol 3, 4, 5-trisphosphate without subsequent activation. Biochemical Journal 1996, 315, 709–713.

(11) Lemmon, M. A. Pleckstrin homology (PH) domains and phosphoinositides. Biochemical Society Symposia. 2007; pp 81–93.

(12) Harlan, J. E.; Hajduk, P. J.; Yoon, H. S.; Fesik, S. W. Pleckstrin homology domains bind to phosphatidylinositol-4, 5-bisphosphate. Nature 1994, 371, 168–170.

(13) Jiang, Z.; Liang, Z.; Shen, B.; Hu, G. Computational analysis of the binding specificities of PH domains. BioMed research international 2015, 2015, 792904.

(14) Carpten, J. D.; Faber, A. L.; Horn, C.; Donoho, G. P.; Briggs, S. L.; Robbins, C. M.; Hostetter, G.; Boguslawski, S.; Moses, T. Y.; Savage, S., et al. A transforming mutation in the pleckstrin homology domain of AKT1 in cancer. Nature 2007, 448, 439–444.

(15) Landgraf, K. E.; Pilling, C.; Falke, J. J. Molecular mechanism of an oncogenic mutation that alters membrane targeting: Glu17Lys modifies the PIP lipid specificity of the AKT1 PH domain. Biochemistry 2008, 47, 12260.

(16) Falkenburger, B. H.; Jensen, J. B.; Dickson, E. J.; Suh, B.-C.; Hille, B. Symposium Review: Phosphoinositides: lipid regulators of membrane proteins. The Journal of physiology 2010, 588, 3179–3185.

(17) Czech, M. P. PIP2 and PIP3: complex roles at the cell surface. Cell 2000, 100, 603–606.

(18) Insall, R. H.; Weiner, O. D. PIP3, PIP2, and cell movement—similar messages, different meanings? Developmental cell 2001, 1, 743–747.

(19) Graber, Z. T.; Thomas, J.; Johnson, E.; Gericke, A.; Kooijman, E. E. Effect of H-bond donor lipids on phosphatidylinositol-3, 4, 5-trisphosphate ionization and clustering. Biophysical Journal 2018, 114, 126–136.

(20) Kooijman, E. E.; King, K. E.; Gangoda, M.; Gericke, A. Ionization properties of phosphatidylinositol polyphosphates in mixed model membranes. Biochemistry 2009, 48, 9360–9371.

(21) Graber, Z. T.; Jiang, Z.; Gericke, A.; Kooijman, E. E. Phosphatidylinositol-4, 5-bisphosphate ionization and domain formation in the presence of lipids with hydrogen bond donor capabilities. Chemistry and Physics of Lipids 2012, 165, 696–704.

(22) Hummer, G.; Köfinger, J. Bayesian ensemble refinement by replica simulations and reweighting. The Journal of chemical physics 2015, 143.

(23) Bottaro, S.; Bengtsen, T.; Lindorff-Larsen, K. Structural bioinformatics: methods and protocols; Springer, 2020; pp 219–240.

(24) Hu, H.; Lu, Z.; Yang, W. Fitting molecular electrostatic potentials from quantum mechanical calculations. Journal of chemical theory and computation 2007, 3, 1004–1013.

(25) Thomas, C. C.; Deak, M.; Alessi, D. R.; van Aalten, D. M. High-resolution structure of the pleckstrin homology domain of protein kinase b/akt bound to phosphatidylinositol (3, 4, 5)-trisphosphate. Current Biology 2002, 12, 1256–1262.

(26) Huang, B. X.; Akbar, M.; Kevala, K.; Kim, H.-Y. Phosphatidylserine is a critical modulator for Akt activation. Journal of Cell Biology 2011, 192, 979–992.

(27) Klauda, J. B.; Venable, R. M.; Freites, J. A.; O’Connor, J. W.; Tobias, D. J.; Mondragon-Ramirez, C.; Vorobyov, I.; MacKerell Jr, A. D.; Pastor, R. W. Update of the CHARMM all-atom additive force field for lipids: validation on six lipid types. The journal of physical chemistry B 2010, 114, 7830–7843.

(28) Essmann, U.; Perera, L.; Berkowitz, M. L.; Darden, T.; Lee, H.; Pedersen, L. G. A smooth particle mesh Ewald method. The Journal of chemical physics 1995, 103, 8577– 8593.

(29) Evans, D. J.; Holian, B. L. The nose–hoover thermostat. The Journal of chemical physics 1985, 83, 4069–4074.

(30) Parrinello, M.; Rahman, A. Polymorphic transitions in single crystals: A new molecular dynamics method. Journal of Applied physics 1981, 52, 7182–7190.

(31) Roux, B. The calculation of the potential of mean force using computer simulations. Computer physics communications 1995, 91, 275–282.

(32) Hub, J. S.; De Groot, B. L.; Van Der Spoel, D. g_wham A free weighted histogram analysis implementation including robust error and autocorrelation estimates. Journal of chemical theory and computation 2010, 6, 3713–3720.

(33) Henderson, L. J. Concerning the relationship between the strength of acids and their capacity to preserve neutrality. American Journal of Physiology-Legacy Content 1908, 21, 173–179.

(34) Jose, G. P.; Gopan, S.; Bhattacharyya, S.; Pucadyil, T. J. A facile, sensitive and quantitative membrane-binding assay for proteins. Traffic 2020, 21, 297–305.

(35) Jose, G. P.; Pucadyil, T. J. Plimap: Proximity-Based labeling of Membrane-Associated proteins. Current Protocols in Protein Science 2020, 101, e110.

(36) Schindelin, J.; Arganda-Carreras, I.; Frise, E.; Kaynig, V.; Longair, M.; Pietzsch, T.; Preibisch, S.; Rueden, C.; Saalfeld, S.; Schmid, B., et al. Fiji: an open-source platform for biological-image analysis. Nature methods 2012, 9, 676–682.

(37) Soubias, O.; Pant, S.; Heinrich, F.; Zhang, Y.; Roy, N. S.; Li, J.; Jian, X.; Yohe, M. E.; Randazzo, P. A.; Lösche, M., et al. Membrane surface recognition by the ASAP1 PH domain and consequences for interactions with the small GTPase Arf1. Science Advances 2020, 6, eabd1882.

(38) Baratam, K.; Jha, K.; Srivastava, A. Flexible pivoting of dynamin pleckstrin homology domain catalyzes fission: insights into molecular degrees of freedom. Molecular Biology of the Cell 2021, 32, 1306–1319.

(39) Li, Z.; Venable, R. M.; Rogers, L. A.; Murray, D.; Pastor, R. W. Molecular dynamics simulations of PIP2 and PIP3 in lipid bilayers: determination of ring orientation, and the effects of surface roughness on a Poisson-Boltzmann description. Biophysical journal 2009, 97, 155–163.

(40) Naughton, F. B.; Kalli, A. C.; Sansom, M. S. Association of peripheral membrane proteins with membranes: free energy of binding of GRP1 PH domain with PIP-containing model bilayers. The journal of physical chemistry letters 2016, 7, 1219.

(41) Beauchamp, K. A.; Pande, V. S.; Das, R. Bayesian energy landscape tilting: towards concordant models of molecular ensembles. Biophysical journal 2014, 106, 1381–1390.

(42) Larsen, A. H.; Wang, Y.; Bottaro, S.; Grudinin, S.; Arleth, L.; Lindorff-Larsen, K. Combining molecular dynamics simulations with small-angle X-ray and neutron scattering data to study multi-domain proteins in solution. PLoS computational biology 2020, 16, e1007870.

